# Tubulin transforms Tau and α-synuclein condensates from pathological to physiological

**DOI:** 10.1101/2025.02.27.640500

**Authors:** Lathan Lucas, Phoebe S. Tsoi, My Diem Quan, Kyoung-Jae Choi, Josephine C. Ferreon, Allan Chris M. Ferreon

**Affiliations:** Department of Biochemistry and Molecular Pharmacology, Baylor College of Medicine, Houston, TX, USA

**Keywords:** Tau, α-synuclein, tubulin, microtubules, biomolecular condensates, liquid-liquid phase separation, FLIM, FRET-FLIM, smFRET, protein aggregation, amyloid

## Abstract

Proteins phase-separate to form condensates that partition and concentrate biomolecules into membraneless compartments. These condensates can exhibit dichotomous behaviors in biology by supporting cellular physiology or instigating pathological protein aggregation^1–3^. Tau and α- synuclein (αSyn) are neuronal proteins that form heterotypic (Tau:αSyn) condensates associated with both physiological and pathological processes. Tau and αSyn functionally regulate microtubules^8–12^, but are also known to misfold and co-deposit in aggregates linked to various neurodegenerative diseases^4,5,6,7^, which highlights the paradoxically ambivalent effect of Tau:αSyn condensation in health and disease. Here, we show that tubulin modulates Tau:αSyn condensates by promoting microtubule interactions, competitively inhibiting the formation of homotypic and heterotypic pathological oligomers. In the absence of tubulin, Tau-driven protein condensation accelerates the formation of toxic Tau:αSyn heterodimers and amyloid fibrils. However, tubulin partitioning into Tau:αSyn condensates modulates protein interactions, promotes microtubule polymerization, and prevents Tau and αSyn oligomerization and aggregation. We distinguished distinct Tau and αSyn structural states adopted in tubulin-absent (pathological) and tubulin-rich (physiological) condensates, correlating compact conformations with aggregation and extended conformations with function. Furthermore, using various neuronal cell models, we showed that loss of stable microtubules, which occurs in Alzheimer’s disease and Parkinsons disease patients^13,14^, results in pathological oligomer formation and loss of neurites, and that functional condensation using an inducible optogenetic Tau construct resulted in microtubule stablization. Our results identify that tubulin is a critical modulator in switching Tau:αSyn pathological condensates to physiological, mechanistically relating the loss of stable microtubules with disease progression. Tubulin restoration strategies and Tau-mediated microtubule stabilization can be potential therapies targeting both Tau-specific and Tau/αSyn mixed pathologies.

---

Deposits of Tau and αSyn fibrillary aggregates in neurons are histopathological hallmarks of neurodegenerative diseases (NDs) such as Alzheimer’s disease (AD) and Parkinson’s disease (PD)^7,23–25^. Interactions between Tau and αSyn in NDs through prion-like infectivity, cross-seeding, and cross-fibrillation have been observed in animal models^7,26,27^ and confirmed in human PD patients^28^. Tau and αSyn undergo heterotypic liquid-liquid phase separation (LLPS) through dynamic interactions within condensates that facilitate pathological aggregate formation^1,20–22,34,46^. Although preventing the formation of these protein-rich condensates is a potential therapeutic approach for NDs, such condensates also play physiological roles and their disruption could impair normal neuronal function^32,47,48,49,50^. For example, Tau-rich condensates *in vitro* and in cells recruit tubulin and provide prime environments for microtubule assembly^32,48^, while αSyn, typically recruited into condensates by proteins such as synapsin, function to regulate synaptic vesicles^49,50^. Interestingly, both Tau and αSyn have functional roles in microtubule regulation: Tau is a well-known microtubule stabilizer^18^ while αSyn modulates microtubule assembly^10–12,19^. Tau and αSyn function cooperatively during brain development^9^ and organize microtubule-dependent transport in neuronal dendrites^8^. Thus, here, we investigated the interactions among Tau, αSyn, and tubulin within condensates to determine if their physiological roles in microtubule regulation could complete with and hinder pathological aggregation.

We tracked protein interactions and morphological evolution within condensates using time-resolved biochemical techniques, combinatorial high-resolution microscopy, and neuronal-based assays to investigate tubulin’s role in modulating and preventing the formation of pathogenic species within condensates. In the absence of tubulin, Tau/αSyn condensates shift to stable amyloidal states, formed via Tau:αSyn heterodimers (TS) and high molecular weight (HMW) oligomers. However, in the presence of tubulin, Tau and αSyn undergo conformational changes, dynamically binding tubulin, assembling microtubule networks, and disfavoring formation of TS heterodimers and HMW oligomers. Our results demonstrate that tubulin acts as a functional scaffold, enabling Tau and αSyn to avoid pathological aggregation. Our findings reveal a novel molecular mechanism, offering a potential therapeutic strategy to selectively inhibit the formation of toxic aggregates while preserving the critical physiological processes required for normal neuronal function.

## Tau LLPS recruits αSyn to form Tau:αSyn pathological condensates

Tau and αSyn are intrinsically disordered proteins (IDPs, Fig. 1a) that, based on *in silico* prediction, exhibit high propensities for LLPS (Fig. 1a, Extended data Fig. 1a). Tau, which has been shown to undergo LLPS facilitated by negatively charged molecules in the presence of crowding agents and low salt concentrations^29,30^, readily forms condensates in physiological conditions (Fig. 1b). In contrast, αSyn does not undergo LLPS under physiological conditions (Fig. 1b) and instead requires high protein concentration (≥200 µM), acidic pH conditions, and prolonged incubation to form condensates^5^. However, in the presence of Tau, αSyn is recruited into heterotypic Tau/αSyn condensates (Fig. 1b). While Tau/αSyn condensation is strictly favored with increasing Tau concentrations, higher αSyn concentrations do not necessarily lead to greater condensation likely primarily due to Tau valancy and binding site availability (Fig 1c).

**Fig. 1.**
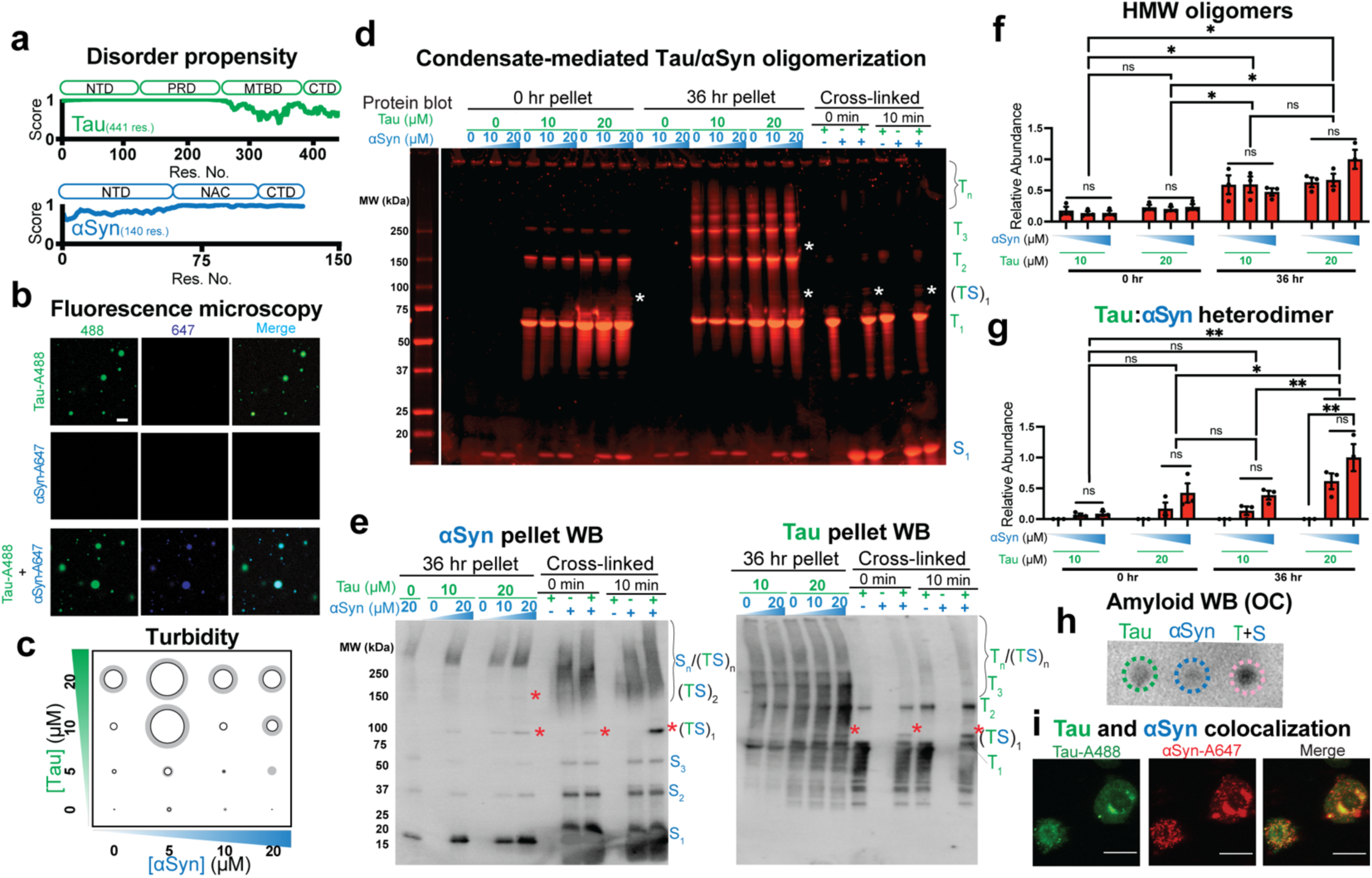
Tau LLPS drives αSyn partitioning and homo- and hetero-oligomerization. **a**, Tau and αSyn domain organization and protein disorder prediction. Both proteins show high disorder propensity (>0.5; scale 0-1, high order to disorder). Tau consists of an N-Terminal Domain (NTD), Proline-Rich Domain (PRD) Microtubule-Binding Domain (MTBD) and C-Terminal Domain (CTD). αSyn consists of an NTD, non-amyloidal component (NAC), and CTD. **b**, Confocal microscopy of 10 µM Tau with ∼200 nM Alexa Fluor 488-labeled Tau (Tau-A488) and/or 10 µM αSyn with ∼200 nM Alexa Fluor 647-labled αSyn (αSyn-A647) mixtures in buffer (10% (v/v) dextran, 5 mM Hepes, pH 8), imaged after 4 hr. Scale bar = 10 µm. **c**, UV absorbance reading at 350 nm of various concentrations of Tau and αSyn mixtures in the same buffer as in ‘**b**’. Circle diameter and shaded region width correspond to the average and SD of 3 replicates. **d**, Sypro Orange-stained- SDS-PAGE gel of Tau/αSyn pellet samples at various concentrations in LLPS buffer (5 mM Hepes, 8 mM Pipes, 10% (v/v) dextran, 0.2 mM MgCl_2_, 0.05 mM EGTA, and 1 mM GTP, pH 7.4) after 0 and 36 hr incubations. Labels on the right of the gels correspond to different oligomeric states (T_n_, Tau monomer/dimer/trimer/oligomers, n = 1,2,3,n; S_n_, αSyn monomer/oligomers, n = 1,n), and white asterisks (*) and TS refer to Tau/αSyn heterodimers. The last 6 lanes correspond to samples crosslinked with 0.8% paraformaldehyde (PFA) for 0 or 10 mins in non-LLPS buffer (PBS) with 20 µM Tau, 80 µM αSyn, or combined. **e**, Tau- (right) and αSyn-specific (left) western blots of a subset of samples shown in **d** showing protein-specific oligomers and the presence of TS when both proteins are present. Samples crosslinked with PFA verified TS with MW sizes that match uncrosslinked samples. **f-g**, Quantitation of gels (represented in **d**) for Tau/αSyn HMW oligomers (bands above monomer in **d**) and TS heterodimers (*), n = 3, statistical analysis: ANOVA multiple comparisons; * p = 0.05, ** p = 0.01, ns = not significant. Error bars = SEM. **h**, Dot blot assay for amyloid detection (anti-OC) with Tau and/or αSyn (20 µM each) in LLPS buffer incubated for ∼3 hours. **i**, N2A cells transfected with 1 µM each of Tau-A488 (green) and αSyn-A647 (red) incubated for 2 days. Puncta shows colocalization of both proteins (yellow). Scale bar = 25 µm.

We next investigated the effects of condensation on Tau:αSyn interactions and on homotypic and heterotypic oligomerization. Tau/αSyn condensates were aged, pelleted, and analyzed using SDS-PAGE to quantify distinct monomeric and oligomeric species (Fig. 1d, Extended Fig. 1b). Within condensates, Tau and αSyn formed SDS-, heat-, and reducing agent-resistant oligomeric species that increased in abundance with time, consistant with previous observations^30,31^ (Fig. 1d; Extended Fig 1b). Western blots of Tau and αSyn verified the presence of stable dimeric, trimeric, and oligomeric species formed within condensates after 36 hr (Fig. 1e). Unexpectedly, exclusively in condensate samples containing both Tau and αSyn, we observed distinct bands corresponding to Tau:αSyn heterodimers (**TS**, denoted with * in Fig. 1d and 1e), which became more evident with longer incubation (72 hr, Extended data Fig. 1c). We confirmed that these bands were indeed TS heterodimers through crosslinking by paraformaldehyde (PFA) in non-LLPS buffer (PBS, Fig. 1e). Crosslinking samples of Tau or high concentrations of αSyn (80 µM) alone produced no TS band, but crosslinking Tau and αSyn mixtures resulted in prominent TS heterodimer bands, matching those observed for Tau/αSyn condensates (Fig. 1d and 1e). All stable Tau/αSyn HMW oligomers significantly increased as a function of time, linking condensate maturation to oligomerization (Fig. 1f). TS heterodimer formation significantly increased with incubation time and also increased proportionally with increasing αSyn concentrations, suggesting that TS heterodimer emergence is primarily dependent on αSyn abundance (Fig 1g). Tau and αSyn are both capable of forming amyloidal structures, but pathogenic amyloid staining was enriched in Tau:αSyn condensates (Fig. 1h). To determine if Tau and αSyn interact within neuroblastoma neuro-2a (N2a) cells, we treated cells with fluorescently labeled Tau and αSyn protein mixtures (Alexa Fluor-labeled Tau-A488 and αSyn- A647), and observed that the proteins colocalized within cytoplasmic puncta (Fig. 1i). Collectively, our data show that LLPS of Tau leads to αSyn recruitment and formation of heterotypic condensates, resulting in an enviorment that favors the emergence of highly stable amyloidogenic hetero- and homotypic oligomers.

## Tubulin transforms Tau/αSyn condensate morphology

Tau condensates recruit and enrich tubulin, surpassing the tubulin critical concentration required for microtubule assembly^29,32^. However, the role of Tau/αSyn condensates in microtubule formation and how tubulin affects TS heterodimer formation have not been studied. To investigate the complex interplay among the three proteins, we generated time-resolved confocal microscopy phase diagrams (Fig. 2a, Extended data Figs. 2-3) to visualize LLPS and microtubule formation across various Tau/αSyn/tubulin concentrations. Tubulin co-partitioned into Tau/αSyn condensates at all concentrations tested. The presence of tubulin transformed condensate shapes from spherical to elongated tactoids: spindle-like organizations of microtubules^33^ (Fig. 2b-c). Formation of microtubule filaments and bundles were more evident with time (Fig. 2a). Fluorescence recovery after photobleaching (FRAP) of Tau/αSyn droplets and Tau/αSyn/tubulin tactoids at 4 hr-incubation showed that both condensates have liquid-like properties consistent with LLPS. Tubulin exhibited slower fluorescence recovery compared to Tau and αSyn, which is consistant with formation of stable microtubule bundles within tactoids (Fig. 2d). From the microscopy phase diagram, we examined partitioning of each protein to condensates and tactoids (Fig. 2e). There were no significant differences in Tau partitioning with varying Tau and αSyn concentrations (10-20 µM) in the absence of tubulin (top left panel), suggesting that dominant role of Tau and passive role of αSyn in the condensation equilibrium. In contrast, we observed αSyn partitioning directly increased as a function of αSyn concentration in Tau/αSyn condensates without tubulin (top middle panel), consistent with greater TS heterodimer formation at higher αSyn levels (Fig. 1d). Similarly, upon addition of tubulin (left lower panels), Tau partitioning generally increased with Tau concentrations wheras αSyn partitioning (middle lower panels) was less dependent on αSyn concentration. Interestingly, we observed that the average Tau and αSyn condensate partitioning decreased as tubulin concentrations increased, which reflect not only the tubulin dependent population redistribution but also condensate droplet to tactoid morphological evolution (Fig. 2e and f). Our observations may also reflect αSyn’s dual microtubule assembly/catastrophe behavior depending on tubulin concentrations^11^. A cartoon morphological diagram (Fig. 2g) summarizes the concentration regimes of tubulin-poor and tubulin-rich Tau/αSyn condensates, and molecular states of the protein components based on cumulative microscopy, gel and amyloid dot blot data. Our findings demonstrate that in the Tau/αSyn/tubulin ternary system, Tau initiates and drives LLPS and recruits αSyn and tubulin, and tubulin regulates dynamic condensate component redistribution and condensate morphological transformation from droplet to tactoid.

**Fig. 2.**
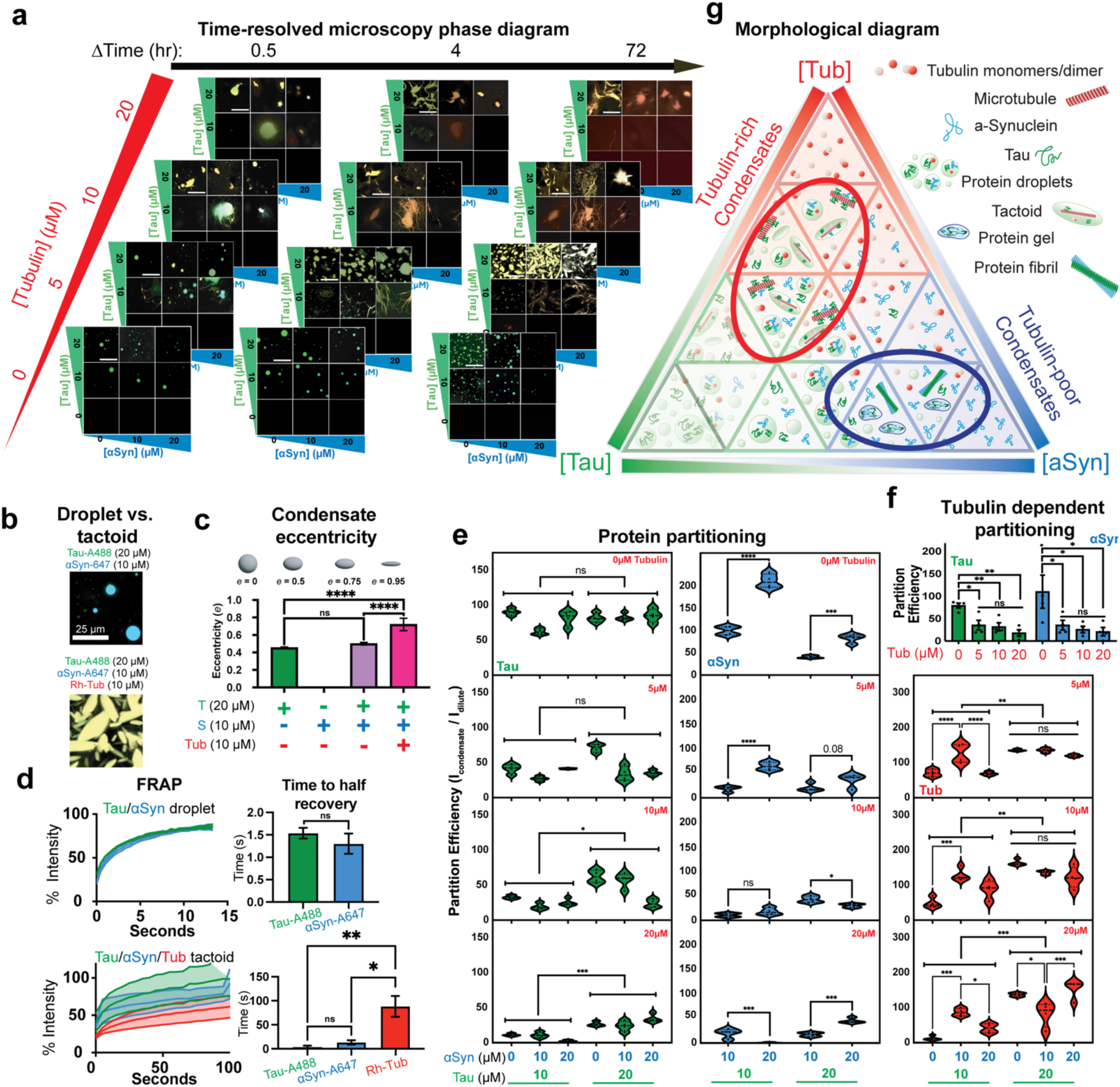
Morphological transformation of Tau and αSyn Condensates in the presence of Tubulin. **a**, Time-resolved (0.5, 4 and 72 hr) confocal microscopy of Tau, αSyn, and Tubulin mixtures at varying concentrations (0-20 µM) in LLPS buffer. Mixtures also included fluorescently labeled Tau-A488, αSyn-A647, and Rhodamine-Tubulin (Rh-Tub) at labeled to unlabeled protein concentration ratios of 1:40, 1:50 and 1:50, respectively. Scale bars = 50 µm. **b**, Representative conditions from **a** showing morphological difference between droplets and tubulin-rich tactoids. **c**, Quantification of shape eccentricity in select conditions (20 µM Tau and 10 µM αSyn ± 10 µM Tubulin after 4 hr). n = 3 images with at least 80 droplets/image. Statistical analysis: one-way ANOVA: **** p = 0.001, ns = not significant. **d**, FRAP data of Tau/αSyn ± Tubulin condensates from **b**. n = 3, Statistical analysis: one-way ANOVA; * p = 0.05, ** p = 0.01, ns = not significant. Error bars = SEM. **e**, Partition efficiency (ratio of mean fluorescence intensity inside condensates vs. in the dilute phase) of Tau/αSyn/Tubulin mixtures (represented in **a**). n = 4, statistical analysis: one-way ANOVA multiple comparisons; * p=0.05, ** p = 0.01, *** p=0.005, **** p=0.001, ns = not significant. **f**, Average partition efficiency (across all Tau and αSyn concentrations in graphs **e**) of Tau (green) and αSyn (blue) at different tubulin concentrations. n = 3, statistical analysis: one-way ANOVA; * p=0.05, ** p = 0.01, ns = not significant. Error bars = SEM. **g**, Cartoon representation of Tau/αSyn/Tubulin ternary system (morphological diagram). The diagram depicts phase and morphological transitions based on confocal images (**a**), western blots of oligomers (**Fig. 1e**) and dot blot amyloid assays (**Fig. 1h**).

## Tubulin prevents formation of pathological Tau/αSyn oligomers and heterodimers

The tubulin-dependent transformation of Tau/αSyn condensates from droplets to tactoids, which actively stabilize microtubules, suggests a dynamic interaction between tubulin, Tau, and αSyn within condensates. We hypothesized that tubulin disrupts homotypic and heterotypic interactions of Tau and αSyn, thereby preventing pathological oligomerization. To test this, we monitored changes in Tau and αSyn oligomerization and TS heterodimer formation in the presence of tubulin using time-resolved western blots (Fig. 3a and b, Extended data Fig. 4a). Because Tau also forms functional dimers and trimers along microtubules^35^, quantitation of HMW Tau oligomers considered only bands corresponding to species larger than trimers. Consistent with Fig. 1d and e data, in the absence of tubulin, Tau/αSyn condensates (pellet fraction) promoted a time-dependent increase in HMW Tau/αSyn homo- and hetero-oligomers (Fig. 3c-e). However within condensates containting tubulin, Tau/αSyn homo-oligomers were significantly reduced, and TS heterodimers were barely visible (Fig. 3b,e), suggesting that tubulin interactions with Tau and αSyn reduce propensities for Tau and αSyn oligomerization and TS heterodimer formation. Without tubulin, Tau binds αSyn’s CTD via its PRD^34^, which is adjacent to the MTBD (Fig. 1a). Tubulin, which interacts with Tau MTBD to stabilize microtubules likely sterically hinders αSyn-Tau interactions and prevents stable TS heterodimer formation. Consistantly, the presence of microtubules, as confirmed by microscopy (Fig. 2a and Extended data Fig. 2 and 3) reduced TS heterodimers (Extended data Fig. 2,3), further demonstrating that tubulin incorporation within Tau/αSyn condensates promotes microtubule formation while reducing pathological aggregation.

**Fig. 3.**
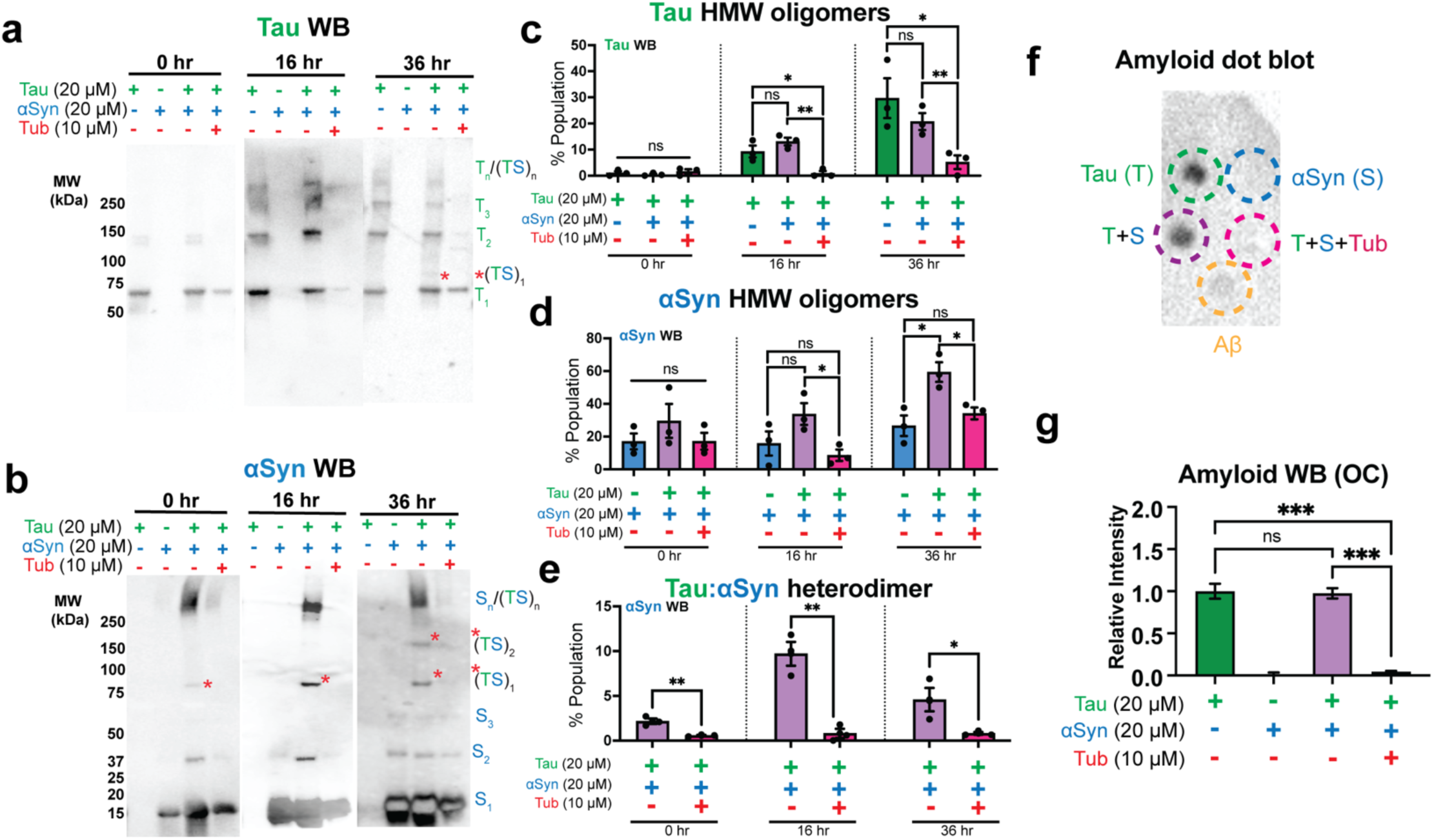
Tubulin prevents formation of Tau/αSyn HMW homo- and hetero-oligomers. **a** and **b,** Representative western blots (anti-Tau, **a**; anti-αSyn, **b**) of Tau/αSyn/Tubulin samples (pellet fraction) incubated for 0, 16, and 36 hr in LLPS buffer. Orange asterisks (*) and *TS refer to Tau/αSyn heterodimers. **c**-**e**, Quantification of bands in blots represented in **a** and **b** (n = 3), showing Tau HMW oligomers (larger than Tau trimer), αSyn HMW oligomers (larger than monomer), and Tau/αSyn (TS) heterodimers, respectively. Statistical analysis: one-way ANOVA multiple comparisons, * p=0.05, ** p = 0.01, ns = not significant. Error bars = SEM. **f**, Representative (n = 3) amyloid dot blot assay (anti-OC) of Tau, αSyn, and tubulin in different combinations (20 µM each), incubated for ∼20 days. 2 µM Amyloid beta (Aβ) was used as reference. **g**, Quantitation of dot blots (n = 3) normalized to Aβ intensity. Statistical analysis: one-way ANOVA multiple comparisons; *** p=0.005, ns = not significant. Error bars = SEM.

The ability of tubulin to outcompete homotypic and heterotypic interactions of Tau and αSyn within condensates suggests that these condensates exhibit detectable Tau:tubulin and/or αSyn:tubulin complexes. αSyn:tubulin oligomers have been previously observed *in vitro*^11^ and in brain homogenates^36^, and Tau is well established to bind tubulin via its MTBD^32,48^. Consistent with this, in Tau:αSyn:tubulin condensates, we detected Western blot bands corresponding to soluble Tau:tubulin and αSyn:tubulin species, which partitioned into the supernatant fraction (Extended data Fig. 4a). These findings align with our microscopy partitioning data (Fig. 2e), further confirming that tubulin prevents Tau/αSyn homo- and heterotypic interactions. The tubulin-dependent reduction in these interactions also correlated with a significant decrease in amyloid formation (Fig. 3g, Extended data Fig. 4b), indicating that tubulin within condensates prevents the pathological transformation of Tau and αSyn. Together, these results demonstrate that tubulin competitively interacts with Tau and αSyn within condensates, functionally promoting microtubule formation while suppressing pathogenic oligomerization.

## Tubulin switches Tau and αSyn conformational states within condensates

The molecular mechanism by which tubulin prevents the formation of amyloidogenic Tau and αSyn oligomers within condensates remains unclear. However, the structural dynamics of Tau and αSyn are intrinsically linked to both their physiological functions and pathological aggregation^20–22^. Therefore, we investigated how tubulin modulates the structural states of Tau and αSyn within condensates. Because Tau and αSyn are IDPs and not amenable to traditional structural techniques, we utilized high-resolution spectroscopy. We used Förster resonance energy transfer (FRET) with site-specific fluorescence labeling to measure dye proximities and assess conformational flexibility in the presence and absence of tubulin. Previous studies indicate that Tau MTBD adopts an extended conformation when bound to microtubules but compacts within amyloid aggregates (Fig. 4a)^15,37,38,43^. To distinguish between these structural states, we flanked the MTBD with donor and acceptor FRET dye-pairs (Fig. 4a) and analyzed Tau’s conformation in both dilute and condensate phases using fluorescence lifetime imaging microscopy (FLIM). In condensates, donor fluorescence lifetimes decreased from ∼3.2 ns in the dilute phase to ∼2.7 ns (Fig. 4b,d), indicating structural compaction consistent with the amyloid structures (Fig. 3f and g, Extended data Fig. 4b). Similarly, αSyn, labeled at positions previously shown to distinguish distinct structural states (Fig. 4a)^20^, exhibited a donor fluorescence lifetime shift from ∼3.4 ns in the dilute phase to ∼2.8 ns in condensates (Fig. 4c,d), indicating compaction again consistant with amyloid formation within condensates. Strikingly, recruitment of tubulin into Tau/αSyn condensates significantly increased donor fluorescence lifetimes for both Tau (∼3.3 ns) and αSyn (∼3.2 ns), resembling values observed in the dilute phases. This suggests that tubulin disfavors pathological structural compaction, instead promoting for extended, physiologically competent conformations (Fig. 4d; Extended Data Figs. 6–8).

**Fig. 4.**
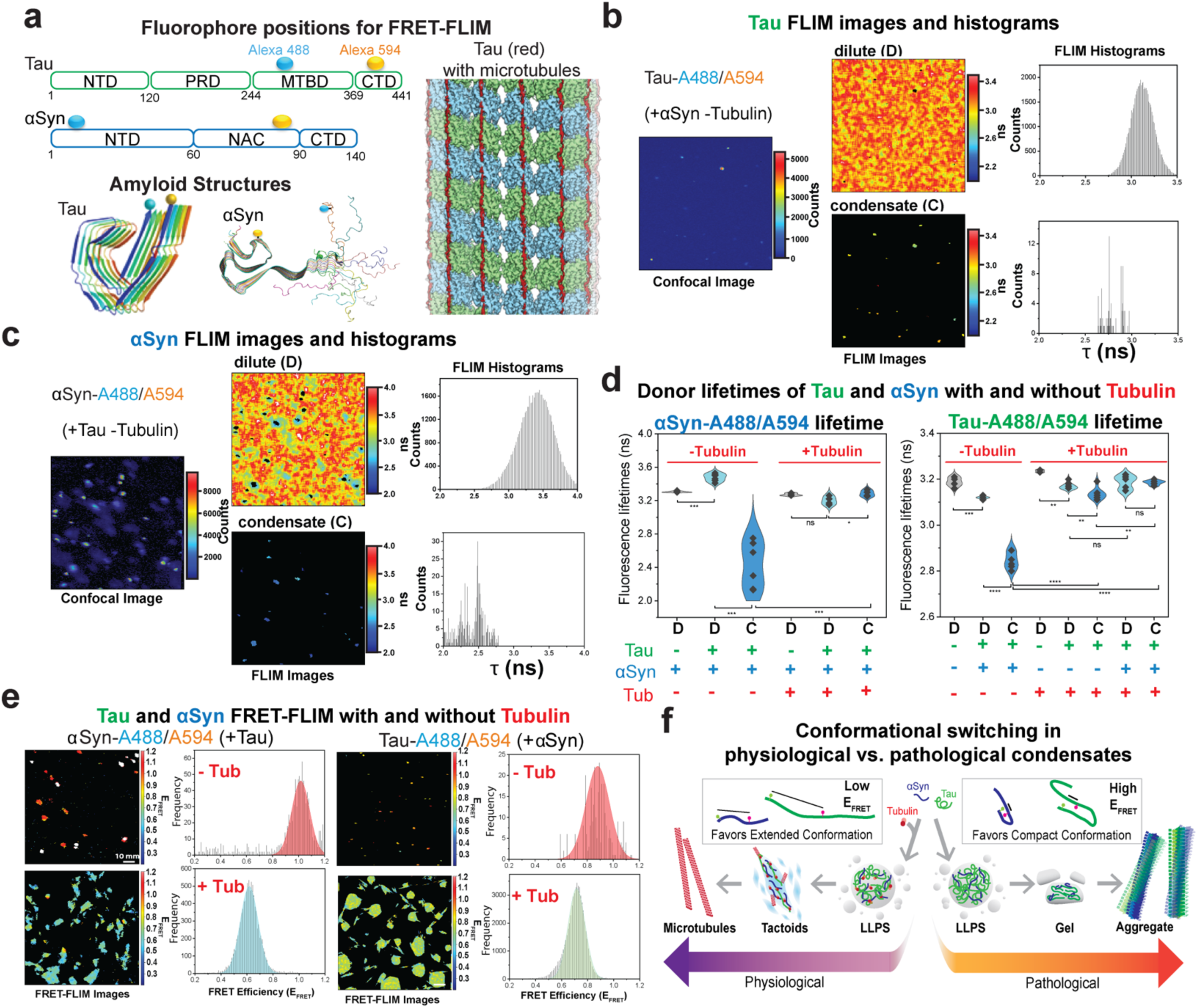
Tau/αSyn conformational switching in tubulin-free and tubulin-rich condensates. **a**, Tau and αSyn domain organization highlighting the locations of the Alexa Fluor dyes (cyan and yellow spheres) for FRET detection. CryoEM structures of Tau amyloids (5O3L), αSyn amyloids (2N0A) and extended Tau on microtubules (EMD-7522). **b**, Representative confocal and FLIM images of Tau (C290/380-A488/594, ∼ 20 nM) in the presence of αSyn, without tubulin. Dilute and condensate images were generated by intensity thresholding, with donor lifetime histograms shown. **c**, Representative confocal and FLIM images of αSyn (C7/84-A488/594, ∼ 20 nM) in the presence of Tau, without tubulin. Dilute and condensate images were generated by intensity thresholding, with donor lifetime histograms shown. **d**, Violin plot of FLIM donor lifetimes of Tau (C291/380- A488/594) (left) or αSyn (C7/84- A488/594) (right) for various conditions (Tau ± αSyn ±Tubulin, Extended Table 1). D and C represent measurements made in the dilute and condensed phases, respectively. n = 5, statistical analysis: One-way ANOVA; * p = 0.05, ** p = 0.01, *** p = 0.001, **** p = 0.0001, ns = not significant. Violin spread = SD, center of violin = mean, width of violin = distribution. **e**, FRET-FLIM images of αSyn (C7/84- A488/594, left) and Tau (C291/380- A488/594, right) in conditions with and without tubulin. FRET-FLIM histograms show distinct αSyn and Tau conformations. **f**, Schematic diagram for how the presence of tubulin transforms Tau/αSyn pathological to Tau/αSyn/tubulin physiological condensates with corresponding favored Tau and αSyn conformations.

To further differentiate structural states in physiological condensates vs. pathological, we converted fluorescence lifetimes to resonance energy transfer efficiencies (E_FRET_), which correlate with inter-dye spacial distances. In the absence of tubulin, Tau and αSyn exhibited compact, amyloid-like states with high E_FRET_ of ∼1 for Tau, and ∼0.9 for αSyn (10–20 Å), consistent with our amyloid-positive dot blot assays (Fig. 1h) and previously reported amyloid structures (Fig. 4a, 4e)^37^. However, in the presence of tubulin or microtubules, both proteins adopted extended conformations with lower E_FRET_ values of ∼0.6 for Tau and ∼0.7 for αSyn (40–50 Å), resembling those observed in physiological Tau-microtubule cryo-EM structures^15^. These findings demonstrate that tubulin facilitates the conformational expansion of αSyn and Tau, shifting condensate charecter from pathological to a physiological states (Fig. 4f).

## Tubulin loss increases Tau/αSyn oligomerization and Tau hyperphosphorylation

The ability for tubulin to prevent Tau and αSyn oligomerization *in vitro* suggests that the loss of tubulin in neuronal cells promote pathological oligomerization of Tau and αSyn. To asses the implications of tubulin loss, we used siRNA to reduce α-tubulin expression by 50% in N2A cells and assessed endogenous Tau/αSyn oligomerization by western blot (Fig. 5a). Although not significant, αSyn oligomerization trends increased with reduced α-tubulin expression. As anticipated, Tau oligomerization significantly increased upon α-tubulin loss. To directly determine if the reduction of α-tubulin in cells resulted in pathological progression, we measured the abundance of hyperphosphorylated Tau oligomers, a form of Tau that is extensively linked to ND pathologies^18^. Loss of α-tubulin led to a more than five-fold increase in hyperphosphorylated HMW Tau oligomers, linking loss of α-tubulin to Tau pathogenesis. Our data highlights a biological mechanism in which tubulin depletion or loss of microtubule stabilization may contribute neurodegenerative cascades.

**Fig. 5.**
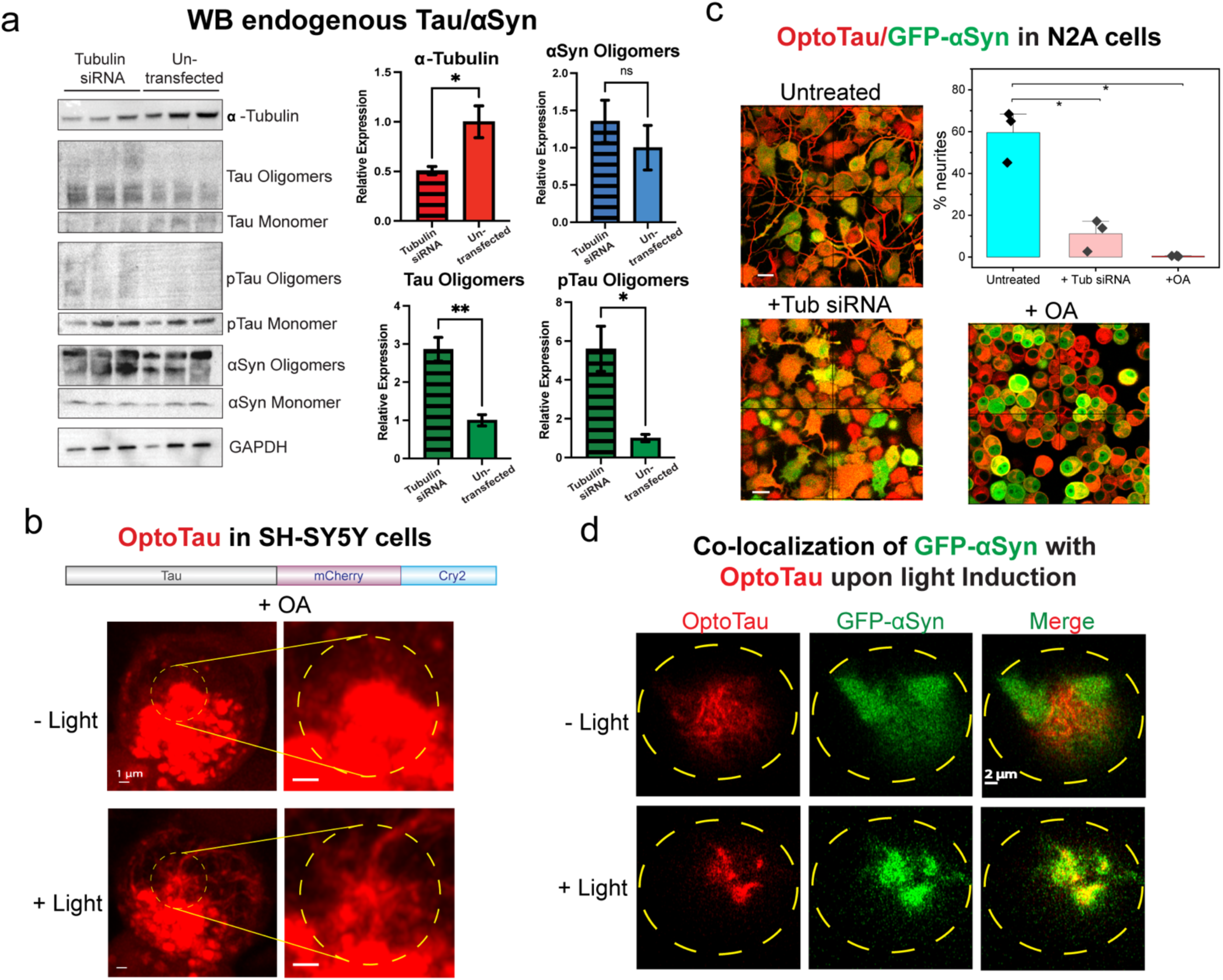
Tubulin knockdown in cells results in toxic oligomers, and microtubule stability is rescuable by OptoTau. **a**, Western blots (anti-α-tubulin, -Tau, -pTau, -αSyn and -GAPDH; top to bottom) of undifferentiated mouse neuroblastoma (N2A) cells with and without tubulin siRNA transfection. Quantification of α-tubulin, αSyn, Tau, and pTau oligomer levels normalized to GAPDH (α-tubulin siRNA transfection, hashed bars vs. untransfected cells, unhashed bars). n = 3, statistical analysis: One-way ANOVA; * p = 0.05, ** p = 0.01, ns = not significant. Error bars = SEM. **b**, Tau optogenetic construct (OptoTau: Tau-mCherry-Cry2, red) stably expressed in undifferentiated human neuroblastoma SH-SY5Y cells. Cells were treated with the oxidative stress agent okadaic acid (OA; 25 nM, 18 hrs). Blue light activation (458 nm laser, 100% power, 20 min) was performed in localized target regions of interest (ROIs) (∼ 5 µm ROI; yellow dash lines). Scale bars = 1 μm. **c**, Confocal microscopy images of N2A cells stably co-expressing OptoTau (red) and GFP-αSyn (green). Cells were either transfected with α-tubulin siRNA or treated with OA (25 nM, 18 hr). Plot of % neurites (ratio of number of neurites vs. total cells, n = 3 biological replicates). Statistical analysis: One-way ANOVA; * p = 0.05. Error bars = SD. Scale bars = 20 µm. **d**, Representative images of N2A cells stably co-expressing OptoTau and GFP-αSyn stressed with 25 nM OA for 18 hr. Cell boundaries are outlined in yellow dashes. Blue light activation (20 min) resulted in enhanced colocalization of OptoTau and GFP-αSyn (yellow). Scale bars = 2 μm.

## OptoTau can rescue microtubule instabilities

Our *in vitro* data (Fig. 2a) and other published data^29,32^ suggest that even with limiting tubulin concentrations, Tau LLPS can facilitate microtubule assembly. To mimic Tau assembly in cells, we designed an optogenetic Tau construct (OptoTau, Fig. 5b) using the blue light-inducible Cry2 dimerization domain^1^. OptoTau was stably expressed in human neuroblastoma SH-SY5Y cells, mostly associated with microtubule filaments. To induce microtubule instabilities, cells were treated with the chemical stressor okadaic acid (OA). Remarkably, blue light activation transformed OptoTau-rich condensates into long, distinct microtubule filaments (Fig. 5b, Extended data Fig. 9). These cell experiments validate our *in vitro* studies and demonstrate the physiological relevance of Tau-rich condensates on supporting microtubule stability.

To assess what happens in the presence of αSyn, we generated N2A cells co-expressing OptoTau and GFP-αSyn (Fig. 5c). As expected, OptoTau colocalizes with microtubules. Although αSyn was shown to colocalize with microtubules in PC12 cells^11^ and rat neurons^10^, the overexpression of GFP-αSyn in our N2A cells resulted in abundant diffuse distribution throughout the nucleus, cytoplasm, as well as in neurite extensions (Fig. 5c). Upon treatment with tubulin siRNA, or OA stress, N2A cells showed a dramatic loss in neurite extensions (Fig. 5c). In addition, cells treated with oxidative stress agents OA and Fe^3+^ (ammonium iron citrate) showed several cells with colocalized Tau/αSyn puncta (Extended data Fig. 10), consistent with co-aggregation of Tau and αSyn. Interestingly, we observed Tau/αSyn punctate colocalization in stressed cells with blue light stimulation (Fig. 5d) further underscoring the dual nature of Tau/αSyn condensates in promoting microtubule stability while also instigating pathological Tau/αSyn co-aggregation under cellular stress.

## Discussion

Our study unveils the role of tubulin in transforming Tau/αSyn pathological condensates to physiological with mechanistic insights into how tubulin influences the structure and behavior of Tau and αSyn within condensates. Tubulin acts as a molecular chaperone, sequestering Tau and αSyn into functional roles in microtubule stabilization and away from aggregation pathways. Moreover, the increased abundance of αSyn and Tau HMW oligomers accompanied by the dramatic increase in hyperphosphorylated Tau oligomers following tubulin depletion in neuronal cells provides compelling evidence that functional interactions with tubulin are protective against Tau and αSyn pathogenic oligomerization and that maintaining tubulin levels and stability above a critical concentration can likely prevent pathological cascade.

In conclusion, our work fundamentally shifts the paradigm of tubulin’s role in neurodegeneration from being merely a structural protein affected by disease to an active protective factor against protein aggregation. This reconceptualization advances two distinct therapeutic avenues. First, while microtubule-stabilizing agents show promise in Alzheimer’s disease (AD) treatment^18,44,45^, our data suggest that Tau-rich condensates that drive microtubule stabilization could also prevent amyloid formation. Second, combining other studies’ findings of reduced tubulin and acetylated tubulin levels in AD and PD^13,14^, with our results that tubulin prevents Tau/αSyn toxic aggregation, we anticipate that tubulin restorative therapy could be a promising therapeutic avenue. Notably, due to the intricate Tau:αSyn interaction network in neurodegenerative pathology, strategies enhancing tubulin and Tau-driven microtubule stability hold promise for targeting mixed Tau:αSyn pathologies at the core of AD and PD.

## ACKNOWLEDGMENTS

We thank the Baylor College of Medicine (BCM) Optical Imaging & Vital Microscopy Core (OIVM) for use of LSM880 confocal microscope. This work was supported by NINDS, NIH grant support to A.C.M.F (R01 NS105874) and Welch Foundation Q-2097-20220331 and NIGMS, NIH grant (R01 GM122763) to J.C.F.

## AUTHOR CONTRIBUTIONS

Conceptualization, A.C.M.F. and J.C.F.; Methodology, L.L., J.C.F. and A.C.M.F.; Software and Formal Analysis, L.L., P.S.T., M.D.Q, K.-J.C., J.C.F. and A.C.M.F.; Investigation, L.L., P.S.T., K.- J.C., M.D.Q., J.C.F., A.C.M.F. and J.C.F.; Visualization, Writing-Original Draft, Review, & Editing: L.L., A.C.M.F. and J.C.F.; Supervision, J.C.F. and A.C.M.F. Funding Acquisition, J.C.F. and A.C.M.F.

## Data availability

Source data for plots, raw data for counts and intensity measurements, and uncropped gel images generated in this study are available from the corresponding authors upon request.

## Code availability

The software used to analyze confocal microscopy, FRET-FLIM data in this study have been listed in the Reporting Summary and are publicly available.

## Statistics & Reproducibility

No statistical method was used to predetermine sample size. No data were excluded from the analyses, and the experiments were not randomized. The Investigators were not blinded to allocation during experiments and outcome assessment.

## Methods

### Bacterial strains

The *E. coli* strains DH5α and BL21 Star (DE3) (Thermo Fisher Scientific, Waltham, MA) were used for plasmid cloning and large-scale preparations of plasmid DNAs, and large-scale protein production, respectively.

### Mammalian cell lines

Lenti-X 293T cells (TaKaRa, Kusatsu, Shiga, Japan) and Neuro-2a (N2A) cells (ATCC, Manassas, VA) were maintained at 37°C, 5% CO_2_ in high glucose DMEM (Corning, Corning, NY) supplemented with 10% heat-inactivated fetal bovine serum (FBS, Corning), 1X MEM Non-Essential Amino Acid Solution (Gibco, Waltham, MA), and 1X Antibiotic-Antimycotic (Gibco). SH- SY5Y cells (ATCC) were maintained at 37°C, 5% CO_2_ in DMEM/F12 (Corning) supplemented with 10% heat-inactivated FBS, 1X MEM Non-Essential Amino Acid Solution, and 1X Antibiotic-Antimycotic. All cells used in this study tested negative for mycoplasma contamination.

### Lentiviral production

Lenti-X 293T (3×10^6^ cells/well) were seeded on a 100 mm diameter cell culture dish and incubated for 1 day at 37°C, 5% CO_2_. The pHR-Tau-mCh-Cry2WT plasmid (10 μg), psPAX2 packaging plasmid (8.4 μg, Addgene #12260, Watertown, MA), and pMD2.G envelope plasmid (2.3 μg, Addgene #12259) were added to 1.5 mL of Opti-MEM medium containing P3000 solution (40 μL, Invitrogen, Walktham, MA). This solution was slowly added to 1.5 mL of Opti-MEM medium containing 2.7% lipofectamine 3000 (Invitrogen) and incubated at room temperature for 20 min. This mixture was then added dropwise to Lenti-X 293T cells in 4 mL of Opti-MEM medium and incubated for 18 hr. The cell culture medium was discarded, and 10 mL of fresh medium (DMEM supplemented with 5% heat-inactivated FBS and 1X MEM Non-Essential Amino Acid Solution) was added and incubated for 1 day. The cell culture medium was collected, and an additional 10 mL of fresh medium was added and incubated for 1 day. The cell culture medium was collected again, and all collected medium was filtered through 0.45 mm-pore size PES filter and concentrated 100-fold using Lenti Concentrator Solution (Origene, Rockville, MD) according to the manufacturer’s instructions. Finally, the lentiviral pellet was resuspended in PBS buffer and either used immediately or snap-frozen and stored at −80°C. The lentiviral titer, determined by the Lentivirus Titer Kit HIV-1 p23 Elisa assay (Origene), was approximately 2-4 x 10^8^ TU/mL.

### Expression of OptoTau (Tau-mCh-Cry2WT) and/or GFP-αSyn in N2A and SH-SY5Y cells

Neuro-2A (N2A,1×10^5^ cells/well) or SH-SY5Y (2×10^5^ cells/well) cells were seeded on a 24-well plate with DMEM supplemented with 5% heat-inactivated FBS and 1X MEM Non-Essential Amino Acid Solution and incubated for 1 day at 37°C, 5% CO_2_. Ten multiplicities of infection (MOI) of lentivirus and 6 mg/mL of polybrene (MilliporeSigma, Burlington, MA) were added to the cells, and the culture dish was incubated for 24 hr, mixing gently. The medium was replaced with DMEM supplemented with 10% heat-inactivated FBS, 1X MEM Non-Essential Amino Acid Solution, and 1X Antibiotic-Antimycotic. The cells were then further incubated for 2 days at 37°C, 5% CO_2_. Tau-mCh-Cry2WT protein expression was confirmed by mCherry fluorescence using EVOS FL Imaging System (Invitrogen). Stable cell lines were established through multiple rounds of serial passaging of the transduced cells. Stable Tau-expressing N2a cells were transfected with EGFP-α-Synuclein-WT plasmid (Addgene #40822) using the Lipofectamine 3000 Transfection Reagent (Invitrogen) according to the manufacturer’s instructions. Transfected cells were selected for 7 days in complete medium containing 0.8 mg/mL Geneticin (G418). Flow cytometry (BD FACSAria II; BD Biosciences, Franklin Lakes, NJ) was used to isolate cells expressing high levels of both mCherry and EGFP fluorescence.

### Mammalian and bacterial expression constructs

Primers in this study are listed in Extended Data Tables 2 and 3. All generated constructs and mutations were confirmed by DNA sequencing (Eurofins Genomics, Louisville, KY).

#### OptoTau (Tau-mCh-Cry2WT)

The human Tau gene was amplified from the tau/pET29b plasmid (gift from Peter Klein, Addgene #16316) by PCR using Phusion High-Fidelity DNA polymerase (Thermo Fisher Scientific) and the primer sets Tau-1F and Tau-1R. The pHR vector DNA fragment was amplified from the pHR-hnRNPA1C-mCh-Cry2WT plasmid (Addgene #101226, a gift from Clifford Brangwynne) using the primer sets Tau-2F and Tau-2R; Extended Data Table 2). The two amplified DNA fragments were ligated to generate the pHR-Tau-mCh- Cry2WT lentiviral transfer plasmid using Gibson Assembly Master Mix (NEB, Ipswich, MA) according to the manufacturer’s instructions.

#### Tau constructs

WT Tau (Addgene #16316) has intrinsic cysteines at residues 291 and 322. To generate cysteine mutants at different desired positions, an *E. coli* codon-optimized human Tau gene (with mutations G196C, C291A, C322A, and G401C) was commercially synthesized (GeneArt, Regensburg, Germany) and incorporated on a pET303 vector. The synthetic plasmid (Tau C196/401) was then used as a template for subsequent mutagenesis to generate other constructs. The QuikChange Site-Directed Mutagenesis method was employed to revert the Tau amino acid residues at position C196, A291, and C401 to WT residues G196, C291, and G401, respectively. Subsequently, the wild-type residues at positions S241 and S341 were mutated to cysteine to generate Tau (C241/341) construct, and mutation of E380 to cysteine to generate a Tau (C291/380) construct that has WT cysteine at residue 291. PCR reactions were performed according to the manufacturer’s instructions (Agilent, Santa Clara, CA). The primer sets used in the PCRs are listed in Extended Data Table 3.

### Protein expression and purification

#### Wild-type Tau and double Cys mutants (Tau (C241/341) and Tau (C291/380))

Expression and purification of WT Tau and Cys mutants were performed using published protocols^29,53^. Corresponding plasmids were transformed into *E. coli* BL21 star cells and grown at 37°C in Terrific Broth medium containing the antibiotics carbenicillin (for WT Tau) and kanamycin (for Cys mutants). The culture was induced with 1 mM isopropyl β-d-1-thiogalactopyranoside (IPTG) when the optical density at 600 nm (OD_600_) reached between 0.8 and 1.0. The cells were grown overnight at 18°C and harvested by centrifugation at 4000×g for 20 min. Pelleted cells were resuspended in 50 mM NaCl, 5 mM DTT, 50 mM sodium phosphate, pH 6.5, and protease inhibitor cocktail (Xpert Protease Inhibitor, GenDEPOT, Katy, TX), and lysed using a homogenizer (Avestin, Ontario, Canada). The lysate was centrifuged at 4°C for 30 min at 50,000×g, and additional salt (to a final concentration of 450 mM NaCl) was added to the supernatant before the solution was incubated for 30 min in near-boiling water (∼85-95°C). The heated solution was centrifuged at 4°C for 2 hrs at 50,000×g and the resulting supernatant was concentrated and subsequently diluted to a final salt concentration of 50 mM NaCl before FPLC (Bio-Rad, Hercules, CA) purification using a heparin sepharose HP column (GE, Chicago, IL) and elution via salt gradient (0-600 mM NaCl). Fractions containing Tau protein were collected, concentrated, and further purified using reverse-phase HPLC (C3 column, Agilent). Fractions containing the final purified protein were collected, aliquoted, lyophilized, and stored at −80°C until later use.

#### α-Synuclein (αSyn)

Expression and purification of αSyn were performed using published protocols^54,55^. Briefly, modified pET-41a (Novagen, EMD Biosciencees, Inc., Gibbstown, NJ) plasmid encoding αSyn protein was transformed into *E. coli* BL21 cells and grown in LB media at 37°C and induced with 1 mM IPTG when OD_600_ reached ∼0.5-0.7. Cells were grown for an additional ∼3 hrs, harvested by centrifugation, and resuspended in 1 mM EDTA, 50 mM NaCl, 50 mM Tris-HCl, pH 7.5. This mixture was frozen and thawed three times by alternating between −80°C and room temperature before sonication and centrifugation. The lysate was incubated at 80°C for ten minutes and centrifuged. The supernatant was further purified using a Q-sepharose (Amersham Biosciences, Amersham, UK) column and eluted using a salt gradient. Fractions containing αSyn were then mixed at a 1:5 ratio (v/v) with a 50 mM sodium acetate solution and added to SP-sepharose (Amersham Biosciences) beads. The pH of the mixture was adjusted to 5.7 and incubated while stirring at 4°C for 1 hr before filtration to remove the beads. The supernatant was then mixed with Q-sepharose beads, the pH adjusted to 4.4, and incubated at 4°C while stirring for 1 hr. The supernatant was once again collected and purified via reverse phase chromatography using a C4 column (Agilent). Fractions containing αSyn were collected, aliquoted, lyophilized, and stored at −80°C until later use.

### Protein sample preparations

Lyophilized Tau and αSyn protein stocks were dissolved in water and centrifuged at 20,000xg for 10 min. Protein samples were filtered through a 0.02 µm filter and protein concentrations were determined (see below). Lyophilized porcine tubulin (Cytoskeleton, Inc., Denver, CO) was dissolved in General Tubulin buffer (Cytoskeleton, Inc.), and snap frozen at −80°C in aliquots. Tubulin was freshly thawed prior to each experiment.

#### Protein concentration determination

The concentrations of recombinant Tau, aSyn, and amyloid beta (Ab, Anaspec, Fremont, CA) proteins were calculated from their UV absorbance extinction coefficients at 280 nm (calculated using Tyr and the Edelhoch method^51,52^; extinction coefficients used are listed in Supplemental Methods). Concentrations of fluorecently labeled proteins were determined using the extinction coefficients of Alexa Fluor 488, 594 and 647 dyes, as appropriate. Tubulin concentrations were determined based on the manufacturer’s instructions (Cytoskeleton, Inc), assuming a molecular weight of 100 kDa for the tubulin heterodimer.

### Preparation of fluorescently labeled proteins

For singly fluorescently labeled samples, purified Tau (WT and Cys mutants) and αSyn were dye-labeled using N-hydroxysuccinimide (NHS) ester chemistries with Alexa Fluor 488 (A488) and Alexa Fluor 647 (A647; Invitrogen), respectively. Labeling reactions were conducted at a 1:10 protein:dye ratio and incubated at RT for 1-3 hr, similar to previously described methods^39,20^. For dual fluorescently labeled samples (*i.e*, for FRET-FLIM experiments), purified Tau and αSyn were dye-labeled using maleimide reactions and conducted at a 1:1 (protein:A488) or 1:3 (protein:A594) ratio. Reactions were incubated at RT for 1 hr to overnight, similar to previously described methods^39,20^. Fractions with different fluorescent labeling ratios (A488:A594) were purified by reverse phase HPLC (Agilent). Samples with optimum fluorescence ratios (*i.e*., a smaller proportion of donor A488 peaks, compared to total peaks (donor and acceptor), were assessed by smFRET measurements and subsequently used for FRET-FLIM experiments.

### Solution turbidity (light scattering)

LLPS was monitored by UV light scattering by measuring UV Absorbance reading at 350 nm using a NanoDrop 2000c Spectrophotometer (Thermo Fisher Scientific). Various solutions containing different concentrations of Tau and αSyn were mixed in LLPS buffer (see below) and samples were incubated for 1 min prior to measurement. 3 independent replicates were collected for each experiment.

### Time-resolved confocal microscopy-based phase diagram

#### Tau (WT)

To generate the three-component concentration phase diagram, different solution conditions with varying Tau, αSyn, and Tubulin concentrations (0, 10, 20 µM for Tau and αSyn; 0, 5, 10, 20 µM for Tubulin) were prepared in LLPS buffer (10% (v/v) Dextran, 0.2 mM MgCl_2_, 0.05 mM EGTA, 1 mM GTP, 5 mM Hepes, 8 mM Pipes, pH 7.4). Samples were supplemented with fluorescently labeled proteins at the following labeled to unlabeled ratios: Tau- A488 (1:40), αSyn-A647 (1:50), Rhodamine (Rh)-Tubulin (1:50) (Cytoskeleton, Inc.). Protein solutions were mixed and transferred onto the coverslip of a μ-Dish, 35mm (ibidi, Martinsried, Germany). Each dish was sealed with parafilm to minimize sample evaporation. Samples were incubated for ∼0.5, 4, and 72 hr prior to imaging (Zeiss LSM 880 Confocal Laser Scanning Microscope and equipped with a 40X oil objective lens, Zeiss, Oberkochen, Germany). Protein samples were simultaneously excited with 488-,561-, and 633-nm lasers using emission filter wavelengths that were adjusted to minimize signal overlap. Tiled images (3X3) were collected for each condition, at z-position 7 µm from the coverslip surface. Two independent replicates were collected.

#### Tau (C241/341)

Similar methods were conducted as described for Tau (WT) above, except with fewer sample conditions: 0 and 5 µM Tubulin and timepoints at 0.5 hr, 4 hr, and 5 days.

#### Tau (C291/380)

Similar methods were conducted as described for Tau (WT) above, except with fewer sample conditions: 0, 5, and 10 µM Tubulin; 0, 5, and 10 µM for both Tau and αSyn. Imaging was performed using Airyscan and separate excitation by 488/561/633 nm lasers. *Image processing and analysis.* All adjustments (e.g., cropping, color adjustments) and analyses (e.g. intensity measurements) were performed using ImageJ2 v.2.9.0^7^. The eccentricity of condensates was quantified using CellProfiler v.4.2.6^56^.

#### Protein Quantitation

Degrees of protein (Tau-A488/αSyn-A647/Rh-Tubulin) partitioning were calculated based on the ratio of the mean fluorescent intensities measured in the background of the condensates, to fluorescence intensity measured within the condensates (droplets/tactoids/microtubules). Condensate and background images were classified based on intensity thresholds. For simplicity and convenience, measured background intensities were scaled to 1 and measured condensate intensities directly equated to partition efficiency values. Statistical analyses were performed using a t-test (see below), and significant results were plotted in the figures. Bracketed conditions were averaged for statistical analysis.

### Fluorescence Recovery After Photo-bleaching (FRAP) imaging

FRAP imaging of droplets and tactoids were performed at RT using an LSM 880 laser-scanning confocal microscope system with a 40X oil objective lens (Zeiss). Samples were prepared as described above for the confocal microscopy phase diagram. Selected samples (*i.e.*, Tau:aSyn droplets (20:10 mM, respectively) and Tau:aSyn:Tubulin tactoids (20:10:10 mM, respectively)) at the 4 hr incubation timepoint were chosen for FRAP experiments. Different regions of interest (ROI; n=4) were selected for bleaching, and reference ROIs (without bleaching) were drawn in adjacent regions (n=2). Following 2-3 baseline images, ROIs were bleached at 100% laser power (488-, 561-, and 633-nm lasers) until intensities were reduced to 20% of pre-bleaching intensities. Additional images were collected 15-100s post-bleaching to assess fluorescence recovery. The recovery half-times were derived from the time required for the fluorescence signal to reach 50% of maximum recovered intensity.

### Detection of Tau/aSyn/Tubulin HMW oligomers

#### Sypro orange-stained SDS-PAGE gels

To assess the kinetics of Tau/αSyn oligomerization, different solution conditions with varying Tau and αSyn concentrations (0, 10, and 20 µM) were prepared in LLPS buffer (10% (v/v) Dextran, 0.2 mM MgCl_2_, 0.05 mM EGTA, 1 mM GTP, 5mM Hepes, 8 mM Pipes, pH 7.4) and incubated for 0, 16, and 36 hr. Samples were centrifuged at 4°C at 17,000xg for 30 min to separate condensates and high molecular weight (HMW) aggregates from supernatants (dilute phase). Supernatants or pellets were mixed 1:1 (v/v) with 2X SDS loading buffer supplemented with 10 mM DTT and 5 mM TCEP (1X=2% (w/v) SDS, 2 mM DTT, 10% (v/v) glycerol, 12.5 mg bromophenol blue, 20 mM Tris, pH 7). Samples were heated at 95°C for 5 min and loaded onto a 4-20% Criterion TGX Precast Gel (Bio-Rad). The gel was run at 225 V for ∼40 minutes in Tris/Glycine/SDS running buffer (VWR, Radnor, PA). Gels were fixed for 30 minutes in 7.5% acetic acid before staining with 1X Sypro Orange (Invitrogen) for 30 min to 1 hr, and destained with 7.5% acetic acid for 20 sec prior to imaging (excitation filter 530 ± 28 nm and emission filter 605 ± 50 nm; ChemiDoc MP Imaging System (Bio-Rad)).

#### Quantitation

Sypro orange-stained gels were run in independent triplicates. Quantification of the gels was performed using Image Studio Software v 5.2.5 (LI-COR, Lincoln, NE). HMW oligomers constituted any bands that appeared above the molecular weight corresponding to αSyn and Tau monomers. Tau and αSyn heterodimers were identified as the doublet band that matched the molecular weight of the heterodimer. The intensity of the quantified bands was divided by the total intensity of the corresponding lane. All lanes were then normalized to a single reference lane. All statistical analyses between conditions were evaluated using a t-test (see below).

#### Fluorescently labeled samples

To assess Tau/αSyn oligomerization states of fluorescently labeled samples, small aliquots (5 µL) were obtained from the same solutions prepared for the confocal microscopy phase diagram and run on SDS-PAGE gels (4-20% Criterion TGX Precast Gels, Bio-Rad), following standard protocols described above. Gels were imaged using appropriate filters for Tau-A488, αSyn-A647, and Rh-tubulin (ChemiDoc MP Imaging System, Bio-Rad).

### Tau-, aSyn-, and Tubulin-specific western blots

#### Western blot assay

SDS-PAGE samples were incubated at 95°C for 5 min and loaded onto 4-20% Criterion TGX Precast Gels (Bio-Rad). The gels were run at 225 V for ∼35-40 min in Tris/Glycine/SDS running buffer (VWR). The proteins were transferred to a methanol-activated Immuno-Blot PVDF Membrane (Bio-Rad) using the Trans-Blot Turbo system (Bio-Rad) at 2.5 A, 25 V, 7 min, or standard 30 min protocol for mini- or midi-gels in Trans-Blot Turbo Transfer Buffer (Bio-Rad). Membranes were blocked for 5 min with EveryBlot Blocking Buffer (Bio-Rad) and incubated with a combination of primary and secondary antibodies for the protein-specific blots (Primary antibodies: anti-αSyn (4D6, Biolegend, San Diego, CA) [1:1000]; anti-Tau (TAU-5, Life Technologies, Carlsbad, CA) [1:800]; anti-b-tubulin (Cell Signaling Technology, Danvers, MA) [1:1000]). All secondary antibodies (anti-mouse and anti-rabbit HRP-conjugated IgG; Cell Signaling Technology) were used at a 1:3000 dilution. After 4-6 hr at RT, or 4°C overnight with light agitation, each blot was washed 3 times with TBST (GenDepot). Membranes were incubated with SuperSignal West Atto Ultimate Sensitivity Chemiluminescent Substrate (Thermo Fisher Scientific) for less than 1 min and imaged using the ChemiDoc MP Imaging System (Bio-Rad).

#### Kinetics of oligomerization

Different solution conditions with varying Tau and αSyn concentrations (0, 10, and 20 µM) and tubulin (0, 10, and 20 µM) were prepared in LLPS buffer (see above). Solutions were incubated at different timepoints (0, 16, or 36 hr) before centrifugation at 4°C at 20,000xg for 30 min. The supernatant was separated from the pellet, and both were mixed 1:1 (v/v) with 2X SDS loading dye buffer (see above).

#### Chemical crosslinking

To assess interactions between Tau and aSyn in non-LLPS conditions (PBS buffer, Corning), chemical crosslinking was performed using 0.8% paraformaldehyde (PFA) and final protein concentrations of 20 µM Tau and/or 80 µM αSyn. The samples were mixed with 5X SDS loading dye (see above) to stop the reaction at different timepoints (∼0 and 10 min). Tau- and αSyn-specific western blots were performed as described above.

#### Quantitation

All western blots described above were performed in triplicate and quantified using the Image Studio Software v 5.2.5 (LI-COR). The intensities of all quantified bands were normalized to the total intensity in their corresponding lanes. Tau-specific blots were quantified to determine the amount of Tau HMW oligomers (bands above the Tau trimer). αSyn-specific blots were quantified to determine the amount of αSyn HMW oligomers (bands above the αSyn monomer) and Tau:αSyn heterodimers. T-test statistical analysis was performed to assess significance across conditions (see below).

### Amyloid dot blots

#### Sample setup and processing

Solution mixtures with Tau, αSyn, and Tubulin (Cytoskeleton, Inc.) or combinations of the three were prepared by diluting the proteins to their respective concentrations in LLPS buffer. Ab (1-40; Anaspec) was used as a positive control.

Solutions (3 µL) were pipetted onto a cut (∼20x20 mm) PVDF Membrane (Bio-Rad) placed in a μ-Dish, 35 mm (ibidi). Excess buffer was added in corners of the dish and the dish was sealed with parafilm to minimize evaporation. After 19-21 days, the solutions were allowed to evaporate for ∼2 hr. The PVDF membrane was immediately fixed (0.4% paraformaldehyde) for 45 min. After fixation, the blot was washed 3 times with water and blocked for 20-30 min with EveryBlot Blocking Buffer (Bio-Rad). The blot was then incubated with both primary and secondary antibodies (Primary Antibody: Anti-amyloid, (OC, EMD Millipore) [1:1000]; Secondary Antibody: HRP-conjugated Rabbit IgG (Cell Signaling Technology) [1:3000]) overnight at 4°C with agitation. Blots were then washed 3 times with TBST (GenDepot). Membranes were incubated with SuperSignal West Atto Ultimate Sensitivity Chemiluminescent Substrate (Thermo Fisher Scientific) for less than 1 min and imaged using the ChemiDoc MP Imaging System (Bio-Rad).

#### Quantitation

All dot blots described were performed in triplicates and the intensity of the dots were quantified using Image Studio Software v 5.2.5 (LI-COR). T-test statistical analysis was performed for comparisons (see below).

### Measuring endogenous Tau/aSyn aggregation in tubulin knockdown N2A cells

#### Cell culture

N2A (4×10^5^ cell/well) were seeded in triplicates in a 6-well dish (Thermo Fisher Scientific) and incubated at 37°C, 5% CO_2_ overnight. 40 pmol of an siRNA cocktail targeting a-tubulin (sc-29189, Santa Cruz Biotechnology, Dallas, TX) was mixed with Lipofectamine RNAiMAX Transfection Reagent (Thermo Fisher Scientific) and incubated in Opti-MEM (Fisher Scientific, Waltham, MA) for 10 min at RT before direct addition to the cells. The transfection mixture was incubated on the cells for 5 min before the addition of growth media and overnight incubation. Media was then changed and cells were incubated for another day before storage at −80°C.

#### Western blot

Frozen cells were scraped using 1X RIPA buffer (Thermo Fisher Scientific), supplemented with 1X protease inhibitor (GenDepot) and 1X phosphatase inhibitor (GenDepot). Cell solutions were then briefly sonicated for ∼1 min. Lysates were then centrifuged at 15,000xg for 15 min at 4°C. Protein concentrations in the supernatant were determined using the Pierce BCA Protein Assay Kit (Thermo Fisher Scientific). Lysates were diluted and prepared in 4X Laemmli Sample Buffer (Bio-Rad) supplemented with 10% 2-mercaptoethanol to a final protein concentration of 3 µg/µL. Samples were heated to 95°C for 5 min. 30 µg of protein was run on a 4–20% Criterion TGX Stain-Free Precast Protein Gel (Bio-Rad) using Tris/Glycine/SDS Running Buffer (Bio-Rad). The proteins were then transferred to Trans-Blot Turbo Nitrocellulose (Bio-Rad) using the Trans-Blot Turbo system (Bio-Rad) at 2.5 A, 25 V, 7 min in Trans-Blot Turbo Transfer Buffer (Bio-Rad). After blocking for 1 hr at RT with 5% Non-Fat Dry-Milk (Chem Cruz, Dallas, TX) in TBST (GenDEPOT), membranes were blocked for 20-30 min with EveryBlot Blocking Buffer (Bio-Rad) and incubated with their respective primary antibody diluted in blocking buffer (anti- αSyn 4D6 [1:1000], Biolegend; anti-Tau (TAU-5, Life Technology) [1:800]; anti-Phospho-Tau (AT8, Invitrogen), [1:1000]; anti-a-tubulin (11H10, Cell Signaling Technology), [1:1000]) overnight at 4°C with light agitation. The blots were washed 3 times with TBST (GenDEPOT) and incubated with their respective secondary antibody diluted in blocking buffer (Horseradish Peroxidase (HRP)-conjugated Mouse IgG (H&L) Antibody (Jackson Immunoresearch, West Grove, PA, [1:10,000]), HRP-conjugated Rabbit IgG (H&L) Antibody (Jackson Immunoresearch, [1:10,000]) for 1 hr at RT. Membranes were then incubated with 1:10 dilution of substrate (SuperSignal West Atto Ultimate Sensitivity Chemiluminescent Substrate, Thermo Fisher Scientific), and incubated for less than 1 min before imaging (ChemiDoc MP Imaging System, Bio-Rad).

#### Quantitation

Samples were acquired from three biological replicates. Western blots were quantified using the Image Studio™ Lite Software (LI-COR). The intensities of all quantified bands were averaged across all replicates and normalized to the control condition. All statistical analyses between conditions were performed using a t-test in GraphPad Prism v.10.3.0 (see below).

### Transfection of Tau and αSyn fluorescently-labeled proteins into N2A cells

N2A (6×10^4^ cells/well) were seeded on a poly-D-lysine-coated CellView 35/10 mm 4- compartment dish (Greiner Bio-One, Kremsmünster, Austria) and incubated for 1 day. A488- labeled Tau protein (4 μM) and A647-labeled αSyn protein (4 μM) were incubated in Opti-MEM medium containing 1.9% Lipofectamine 3000 at RT for 20 min. The protein mixture was added dropwise, directly to the cells to achieve a final protein concentration of 1 μM each with addition of extra Opti-MEM media. After 18 hr of incubation, the medium was replaced with complete media, and the cells were further incubated for an additional 2 days. Cells were then fixed and imaged by confocal microscopy.

### Tubulin knockdown and oxidative stress in N2A cells

#### Transfection of α-Tubulin siRNA in N2A cells co-expressing Tau and α-Synuclein

N2a (1.5×10^5^ cells/well) were seeded on a μ-Dish, 35mm ibiTreat (ibidi) and incubated for 1 day. The cells were transfected with 30 pmol of α-tubulin siRNA (sc-29189, Santa Cruz Biotechnology) using Lipofectamine RNAiMAX Reagent (9 μL, 0.9% final concentration, Invitrogen) according to the manufacturer’s instructions. Two days post-transfection, the cells were either imaged or used for further experiments.

#### Treatment of cells with okadaic acid (OA) or ammonium iron (III) citrate

N2a cells (1.5×10^5^ cells/well) were seeded on a μ-Dish, 35mm ibiTreat (ibidi) and incubated for 1 day. The cells were treated with 25 nM OA (Cayman Chemicals, Ann Arbor, MI) or 5 mM ammonium iron(III) citrate (Millipore Sigma). After 18 hr of incubation, the cells were imaged using confocal microscopy.

#### Image collection, processing and quantitation of neurites

Confocal images (5x5 tiled images) were collected for 3 biological replicates per cell condition (treated with tubulin siRNA, OA or Fe^3+^). After file conversion of Zeiss- to Nikon-compatible files, the images were analyzed using NIS-Elements software (Nikon, Tokyo, Japan) and the cells were counted using the object count algorithm. The number of neurites were manually counted.

### OptoTau blue light activation in stressed cells

#### Cells

For OptoTau experiments, we utilized human neuroblastoma SH-SY5Y cells stably expressing OptoTau and mouse N2A cells stably expressing both OptoTau and GFP-αSyn. The cells were either transfected with tubulin siRNA (as described above) or stressed with OA (25 nM, 18 hr) prior to blue light activation.

#### Blue light activation

Confocal microscopy 3D z-stacks were collected before and after blue light activation on selected ROIs (∼5 µm diameter). For SH-SY5Y, blue light activation was performed using 100% power, 458-nm laser for ∼20 min. For N2A cells, blue light activation was performed using 100% power, 458-nm laser and 10% power 488-nm laser for ∼20 min.

### FRET-FLIM

Fluorescence lifetime imaging microscopy (FLIM) experiments were performed at RT using a custom-built Alba confocal laser microscopy system (ISS, Champaign, Illinois), which employs Olympus IX81 microscope equipped with an Olympus 60X/1.2 NA water objective lens, galvo-controlled mirrors, and imaging and FastFLIM^TM^ modules (ISS). The instrument was calibrated using rhodamine 110 dye in water (4.0 ns fluorescence lifetime in water) prior to data collection. Tau (C291/380-A488/A594) and αSyn (C7/84-A488/A594) FRET samples were prepared in various solution conditions (Extended Data Table 1). FLIM was analyzed by the frequency domain (FD) lifetime fitting as described previously^40,57,58^. For FD fitting, FLIM images were fitted with binning size of 3x3 pixels to 5 different modulation frequencies. The laser power utilized was typically 5-50 µW. There is no significant dependence between laser power and measured fluorescence lifetimes at the laser powers used. Lifetime values were verified to be independent of binning size or z-slices. Confocal FLIM images were generated using the modules in the VistaVision software V4.2 (ISS). FLIM histograms were generated by VistaVision software V4.2 (ISS) or by re-plotting using the OriginPro software. FRET images were converted from FLIM images using the algorithm in the ISS software, based on the following equation:

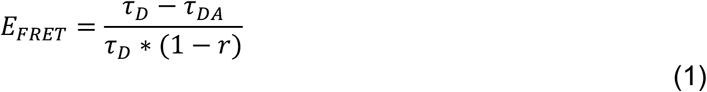

where t_DA_ is the fluorescence lifetime of the donor in the
presence of acceptor (e.g., A488 - A594 FRET labeled sample), t_D_ is the fluorescence lifetime of the donor only (4 ns for A488) and *r* is the unquenched donor peak (ratio of the donor zero peak area to the total peak area in smFRET histograms). For Tau (C291/380-A488/A594) and αSyn (C7/84-A488/A594), *r* values were determined to be 0.71 and 0.66, respectively. The values were back-calculated based on the actual *E_!FRET_* values from smFRET experiments.

### Computational analysis of protein disorder and phase separation propensities

Both human WT Tau (2N4R isoform, uniport ID P10636) and αSyn (uniport ID P37840) protein sequences were used in the analysis. Intrinsically disordered regions were predicted using IUPred2A^59^. Phase separation propensity was predicted using PSPHunter^60^.

### Statistical analysis

Unless otherwise stated in figure legends, student’s t-test was employed for direct comparisons. All *in vitro* gel experiments were performed and quantified in three independent replicates. Image quantification of eccentricity and fluorescent intensity values considered individual condensates and tactoids (n>300 and n>10, respectively) for quantification. Cell culture experiments and subsequent western blots were performed in three biological replicates. Graphs show the average and standard error of measurements (SEM) or standard deviations (SD). Violin plots display the data probability distribution. Figure legends describe the statistical tests employed. Regardless of the analysis method, significance levels were denoted by *, **, ***, and **** for P < 0.05, P < 0.01, P < 0.001, and P < 0.0001. Graphs were generated using either Prism v. 10.3.0 (GraphPad, San Diego, CA) or OriginPro (OriginLab, Northampton, MA).

## Data availability

Source data for plots, raw data for counts and intensity measurements, and uncropped gel images generated in this study are provided in a Source Data file. Any additional information reported in this paper is available from the corresponding authors upon request.

---

**Extended Data Fig. 1.**
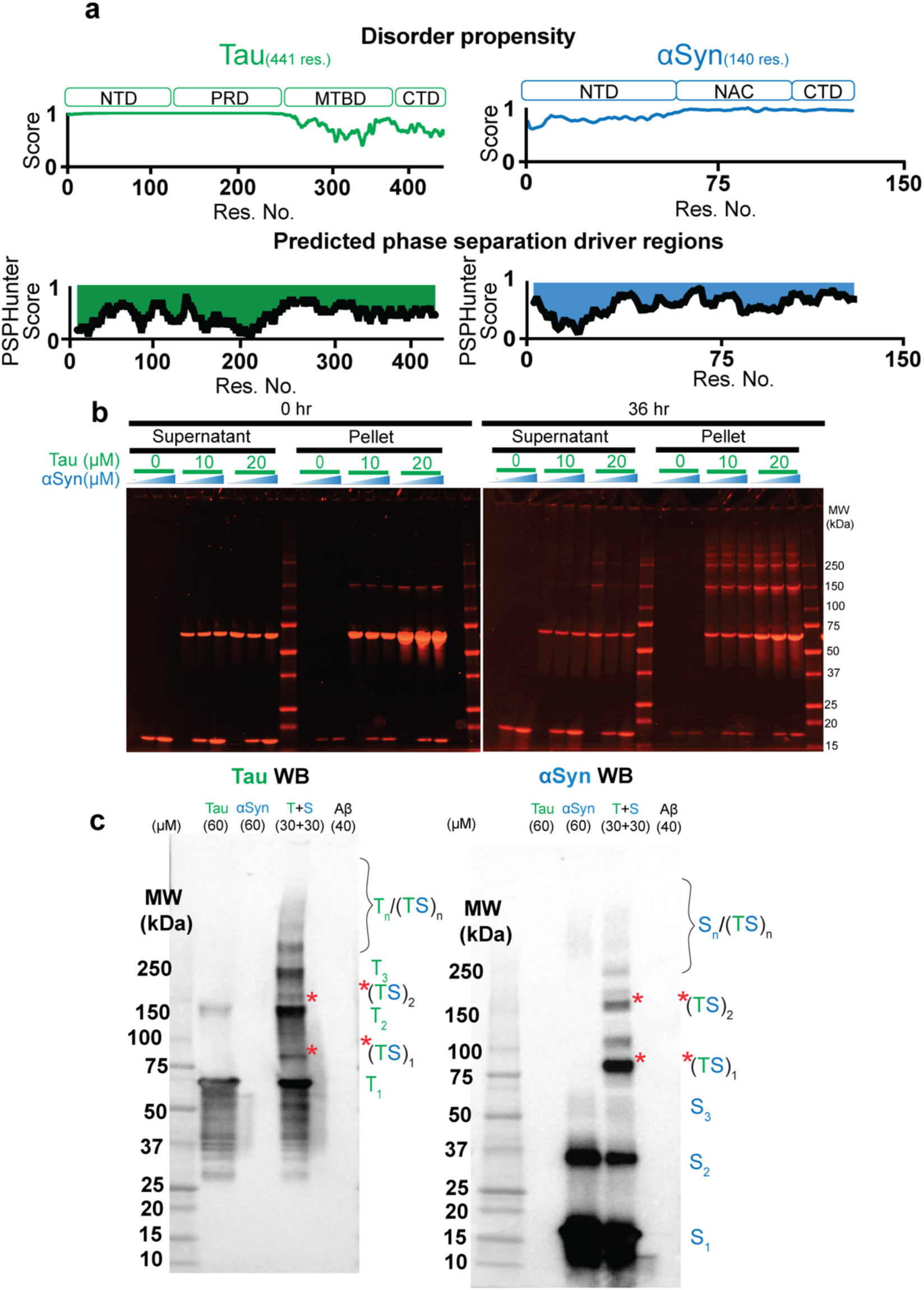
Tau/αSyn phase separation propensities and co-oligomerization. **a**, Protein domain representation (top) of the Tau (green) and αSyn (blue). Tau consists of an N-Terminal Domain (NTD), Proline Rich Domain (PRD), Microtubule-Binding Domain (MTBD), and a C-Terminal Domain (CTD). αSyn consists of an NTD, a non-amyloidal component (NAC), and a CTD. Plots of predicted disorder (middle) where 1 indicates higher disorder and phase separation (bottom) where 0 indicates higher propensities plotted against residue number^2^. **b**, Sypro Orange-stained-SDS-PAGE gels showing different concentration mixtures of Tau/αSyn in LLPS buffer, incubated for 0 and 36 hrs. Samples were centrifuged to separate supernatants and pellets. **c**, Tau- and αSyn-specific western blots of samples at higher concentrations of Tau and αSyn (60 µM individually or 30 µM each when combined; LLPS buffer) aged for 3 days. Aβ (40 µM) was included to check antibody specificity. The blots show the appearance of multiple high molecular weight (HMW) oligomers and Tau:αSyn heterodimer/hetero-oligomers (*). Labels on the right of the gels correspond to different oligomeric states (T_n_, Tau monomer/dimer/trimer/oligomers, n = 1-2-3-n; S_n_, monomer/dimer/oligomers, n = 1-2-3-n; and, orange asterisks (*) and *TS refers to the Tau/αSyn heterodimers).

**Extended Data Fig. 2.**
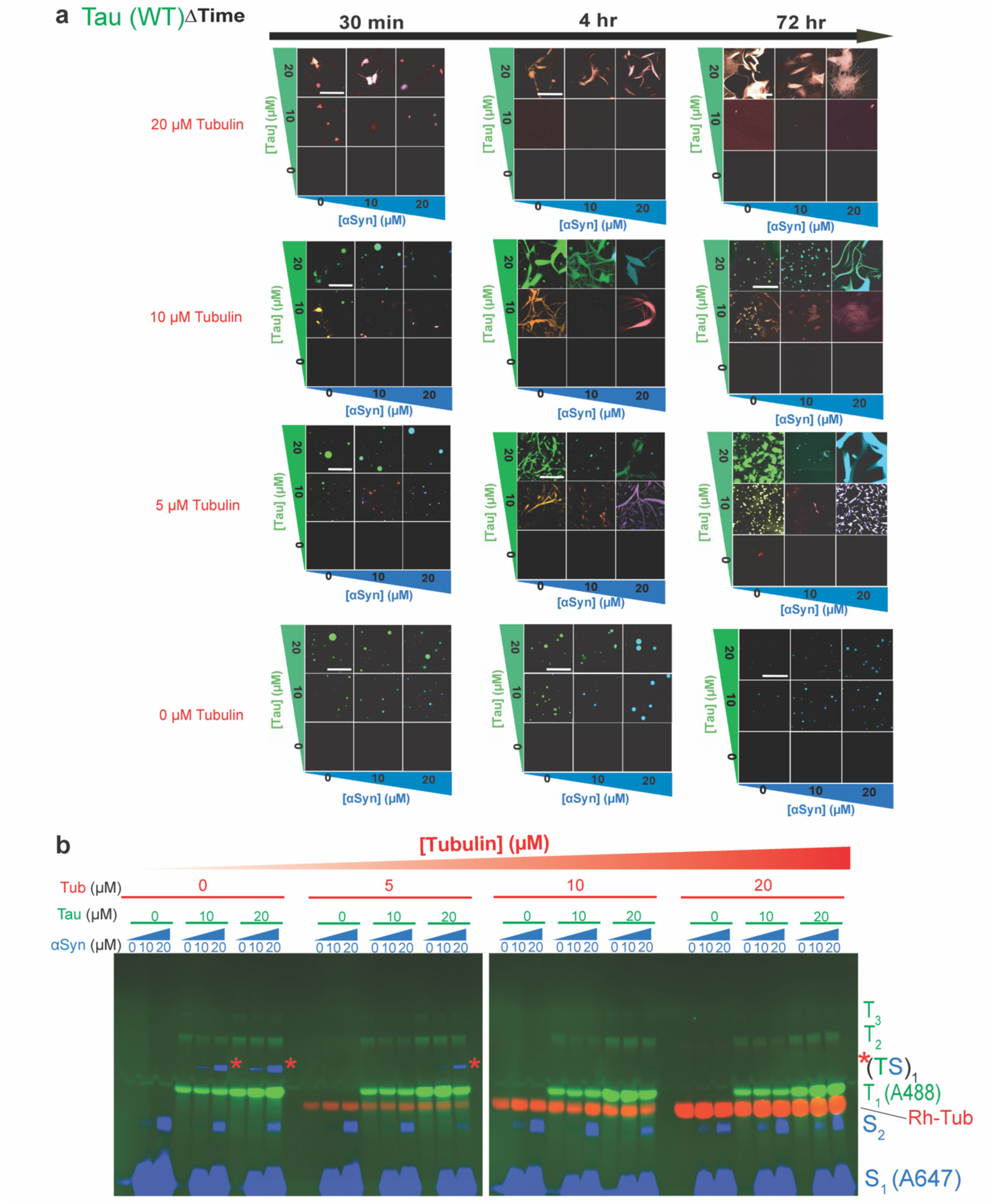
WT Tau/αSyn/Tubulin confocal microscopy phase diagram with corresponding SDS-PAGE gels. **a**, Time-resolved (0.5, 4 and 72 hr) confocal images (independent replicate of Fig **2a** data) at varying concentrations of WT Tau (0, 10 and 20 µM), αSyn (0, 10 and 20 µM), and Tubulin (0, 5, 10 and 20 µM). Fluorescently labeled Tau-A488, αSyn-A647, and Rhodamine-Tubulin (Rh-Tub) were added at labeled to unlabeled ratio of 1:40, 1:50 and 1:50, respectively. Rows with 20 µM Tau (0, 10, 20 µM αSyn) at 5 µM and 10 µM tubulin did not contain Rhodamine-tubulin. Scale bars = 50 µm. **b**, Gels of samples described in **a** at 4 hr incubation. Labels on the right correspond to different oligomeric states (T_1- 3_, Tau monomer to trimer; S_1-2_, monomer/dimer; orange asterisks (*) and *TS refers to the Tau/αSyn heterodimer).

**Extended Data Fig. 3.**
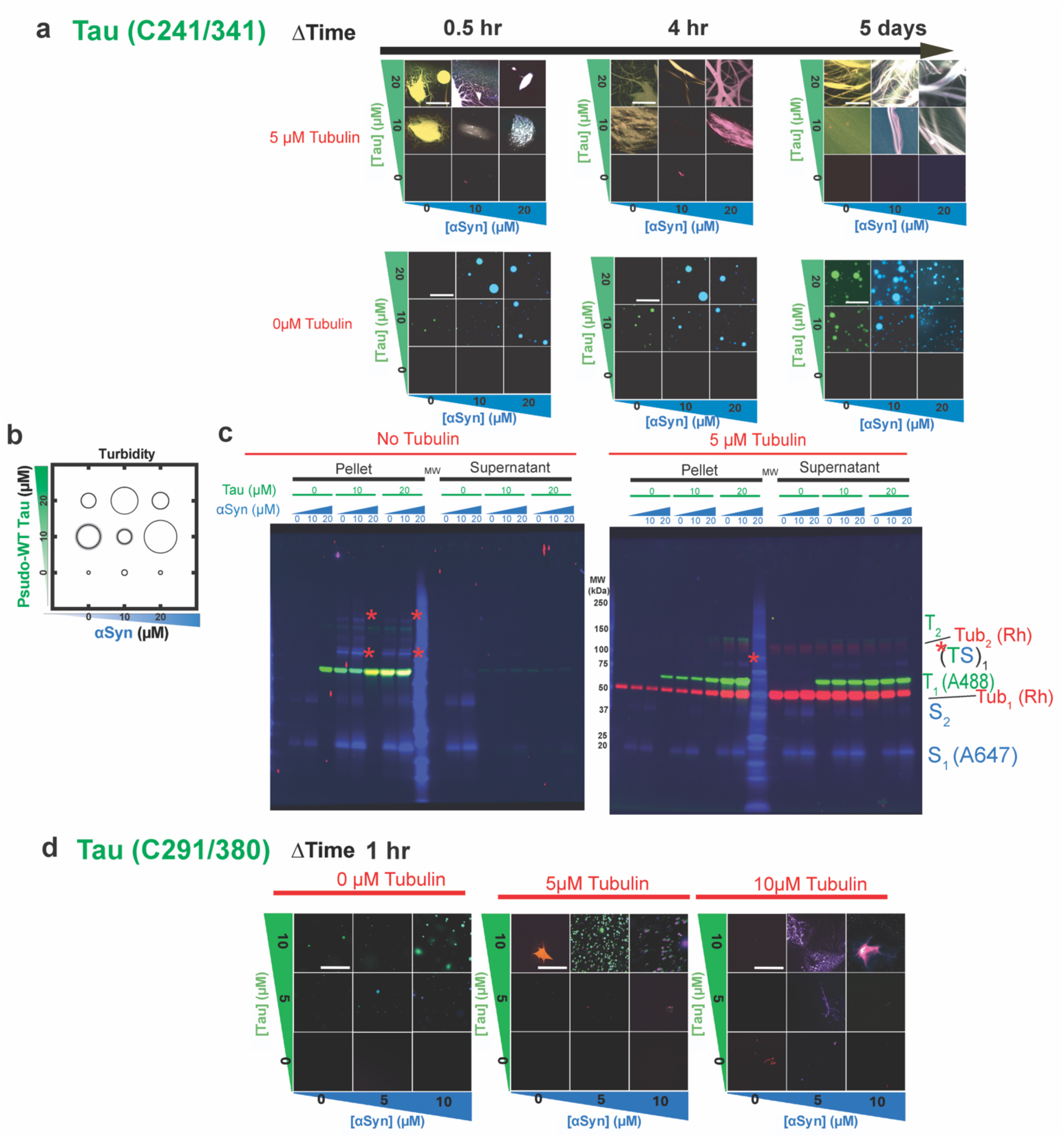
Tau cys mutants/αSyn/Tubulin confocal microscopy phase diagram with corresponding SDS-PAGE gels. **a**, Time-resolved (0.5, 4 and 72 hr) confocal images at varying concentrations of Tau (C241/341); (0, 10 and 20 µM), αSyn (0, 10 and 20 µM) and Tubulin (0 and 5 µM). Fluorescently labeled Tau-A488, αSyn-A647, and Rhodamine-Tubulin (Rh-Tub) were added at labeled to unlabeled ratio of 1:40, 1:50 and 1:50, respectively. Scale bar = 50 µm. **b**, Turbidity based Tau/αSyn phase diagram. **c**, Gels of samples described in **a** at 4 hr incubation. Labels on the right correspond to different oligomeric states (T_1-3_, Tau monomer to trimer; S_1-2_, monomer/dimer; Tub_1-2_, Tubulin monomer/dimer; orange asterisks (*) and *TS refers to the Tau/αSyn heterodimers). **d**, Confocal images of mixtures at varying concentrations of Tau (C291/380); (0, 10 and 20 µM), αSyn (0, 10 and 20 µM) and Tubulin (0 and 5 µM). Scale bar = 50 µm.

**Extended Data Fig. 4.**
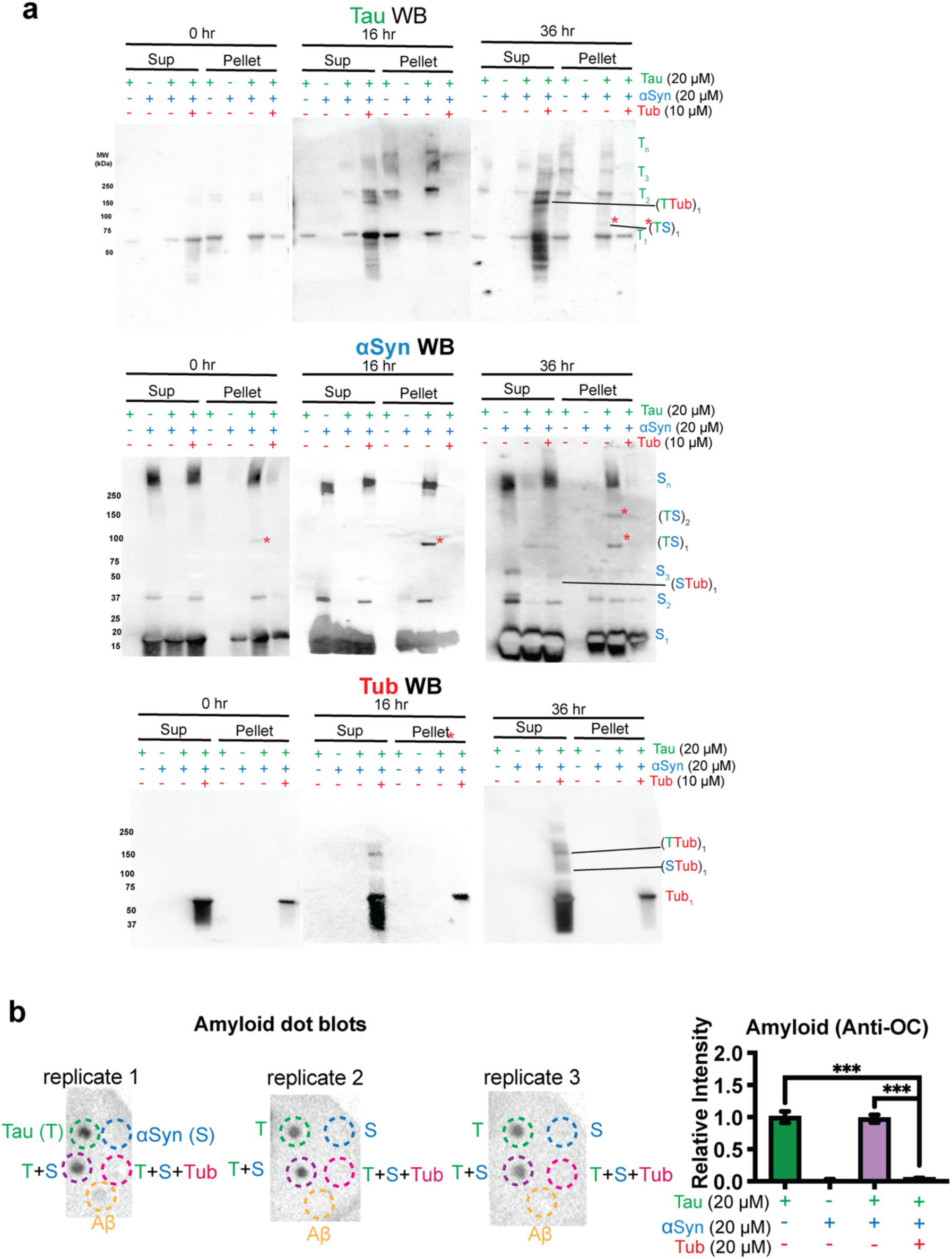
Tubulin prevents Tau/αSyn homo- and hetero-oligomerization and amyloid formation. **a**, Protein-specific western blots of supernatants and pellets of Tau, αSyn or Tubulin samples (top to bottom) incubated at 0, 16 and 36 hr. T_n_, Tau monomer/dimer/trimer/oligomers, n = 1-2-3-n; S_n_, monomer/dimer/trimer/oligomers; n = 1-2-3-n; (T/STub), Tubulin interactions with Tau and αSyn. **b**, Dot blots using an anti-OC antibody to detect amyloids in samples containing Tau or αSyn alone (20 µM each), or Tau and αSyn (20 µM each) in the presence and absence of tubulin (20 µM). Aβ (2 µM) was included as reference. Quantitation is shown in the right panel. Statistical analysis: one-way ANOVA multiple comparisons; *** p=0.005, Error bars = SEM.

**Extended Data Fig. 5.**
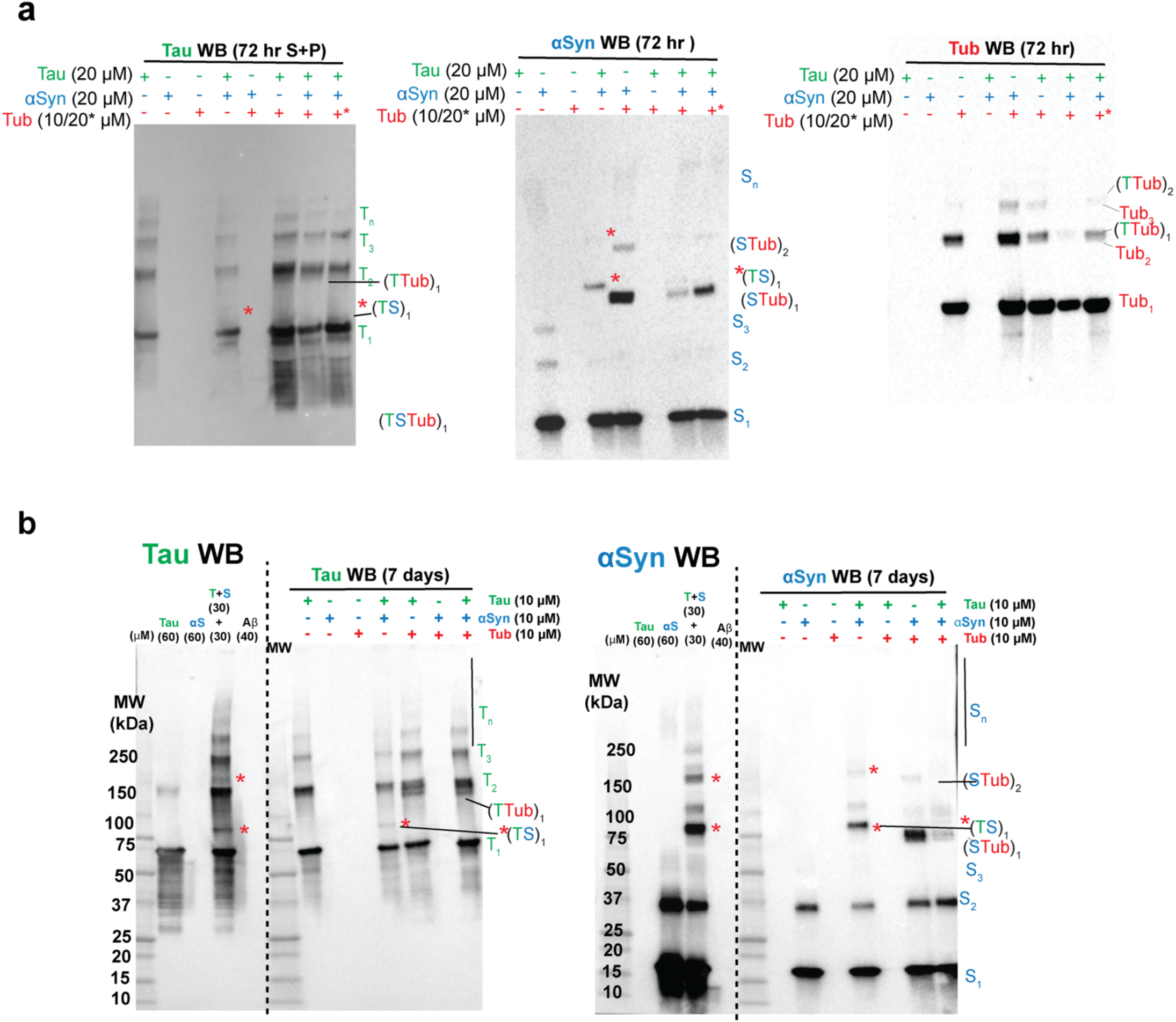
Tubulin prevents Tau/αSyn homo- and hetero-oligomerization. **a**, Protein-specific western blots (Tau, αSyn, Tubulin; left to right, respectively) of 72 hr samples with different combinations of Tau, αSyn, and Tubulin (20, 20 and 10/20 µM, respectively) in LLPS buffer. T_n_, Tau monomer/dimer/trimer/oligomers, n = 1-2-3-n; S_n_, monomer/dimer/trimer/oligomers; n = 1-2-3-n; (T/STub)_n_, Tubulin interactions with Tau and αSyn, n =1-2. **b**, Protein-specific western blots (Tau, αSyn; left to right). The first lanes of the blots correspond to either the moleculae weight marker or samples at higher concentrations of Tau and αSyn (60 µM individually, or 30 µM each when combined; LLPS buffer) aged for 3 days. Aβ (40 µM) was included to check antibody specificity. The last 7 lanes correspond to similarly prepared samples as in **a** but at 10 µM concentration each of Tau, αSyn and/or Tubulin, incubated for 7 days. Labels on the right of the gels correspond to different oligomeric states (T_n_, Tau monomer/dimer/trimer/oligomers, n = 1-2-3-n; S_n_, monomer/dimer/oligomers, n = 1-2-n; orange asterisks (*) and *TS refers to the Tau/αSyn heterodimers; STub refers to αSyn/Tub heterodimer(s) and TTub refer to Tau/Tub heterodimer(s)).

**Extended Data Fig. 6.**
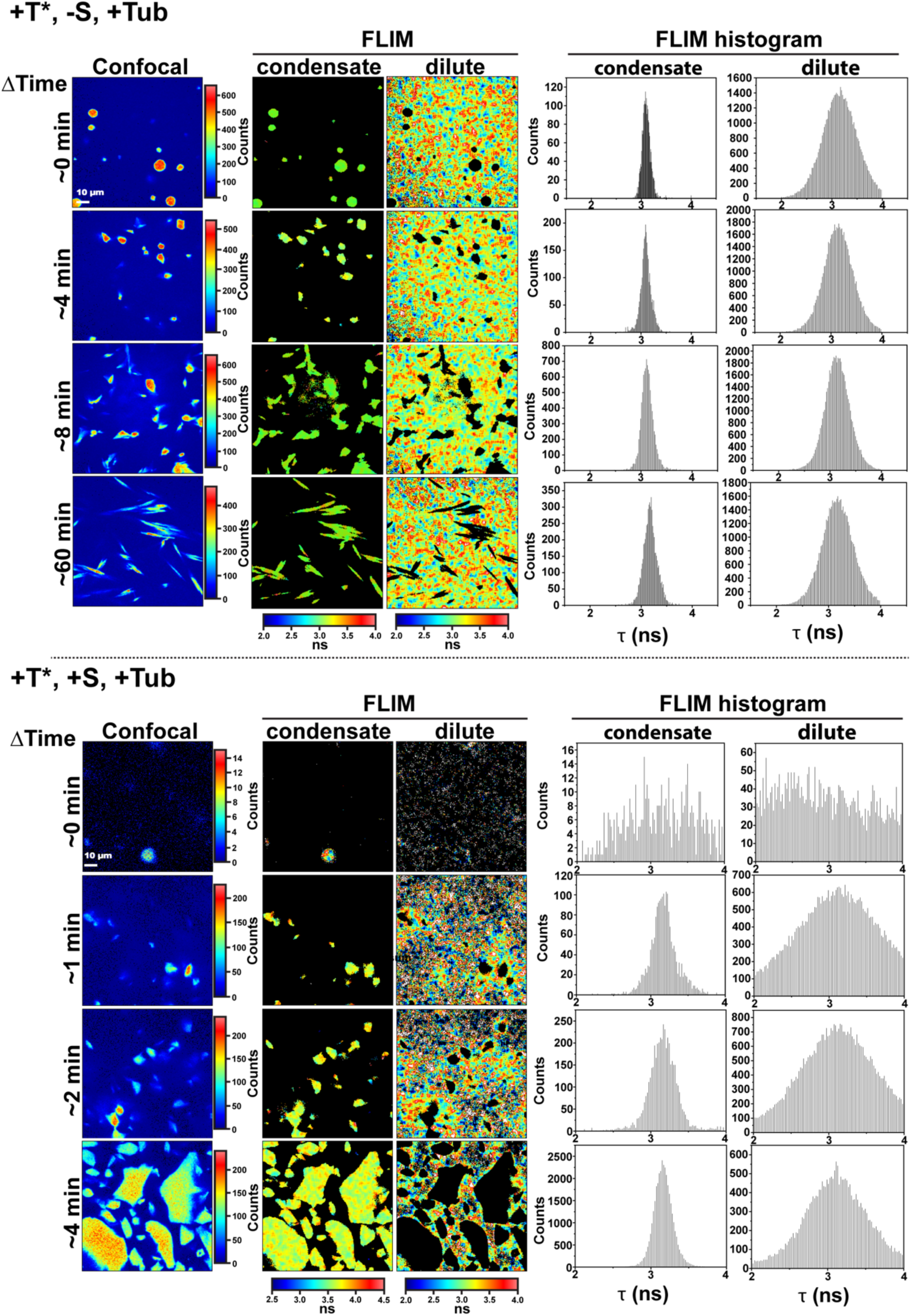
Morphological evolution of Tau/Tubulin Droplets with and without αSyn. Representative confocal and FLIM images of fluorescently labeled 1 nM Tau (A488/A594, T*) with corresponding histograms in the presence of unlabeled 50 µM Tau and 10 µM tubulin, with or without 50 µM αSyn (+T*, -S, +Tub, or +T*, +S, +Tub, respectively) in FRET-FLIM buffer (5 mM Hepes, 8 mM Pipes, 0.2 mM MgCl_2_, 0.05 mM EGTA, and 1 mM GTP, pH 7.4). Separation of dilute vs condensed phases was achieved using intensity thresholding.

**Extended Data Fig. 7.**
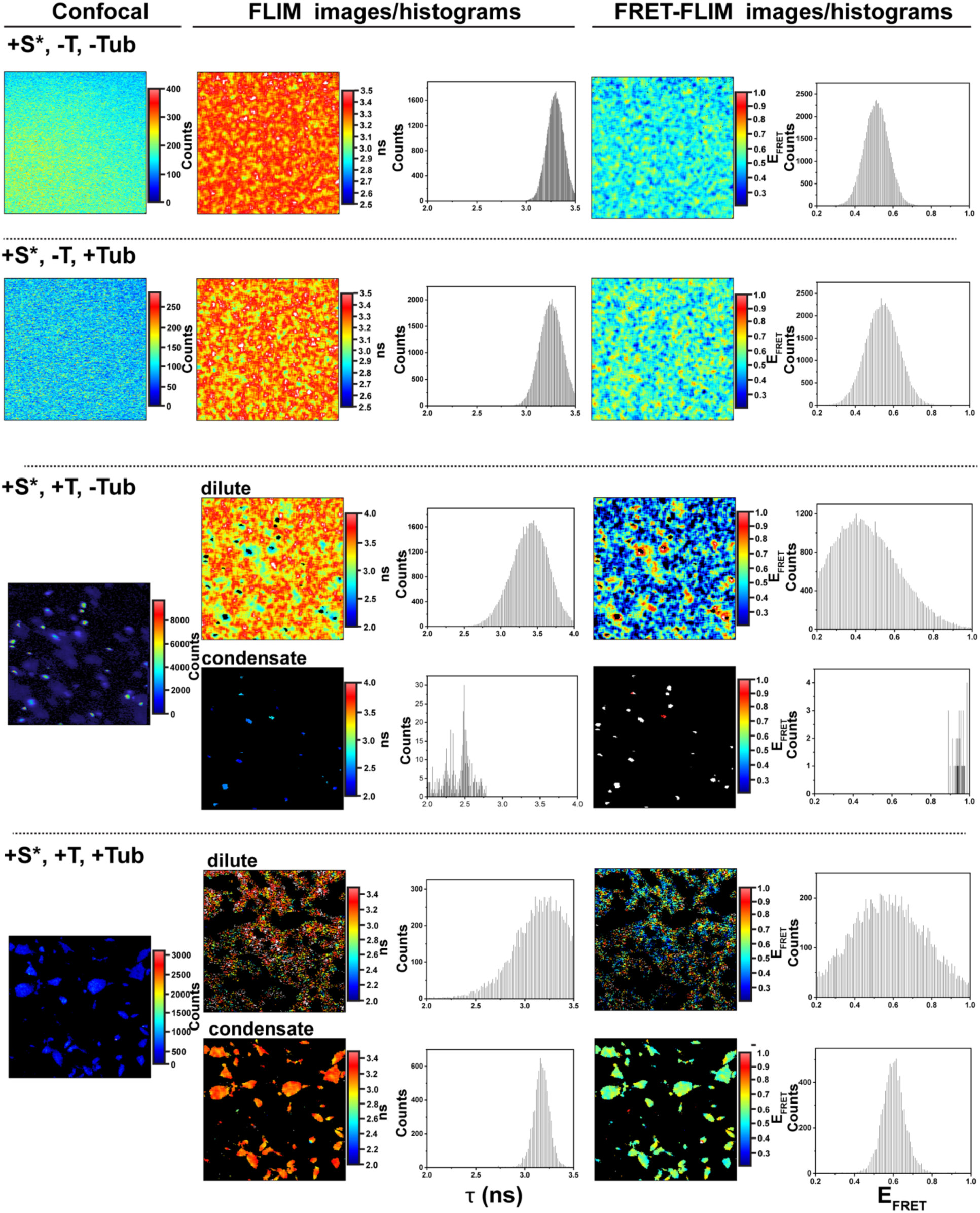
αSyn conformational states modulated by Tau and Tubulin. Representative confocal, FLIM, and FRET-FLIM images with corresponding histograms of fluorescently labeled 1 nM αSyn (A488/A594, S*) in the presence 50 µM unlabeled αSyn 50 µM Tau and/or 10 µM tubulin in FRET-FLIM buffer (5 mM Hepes, 8 mM Pipes, 0.2 mM MgCl_2_, 0.05 mM EGTA, and 1 mM GTP, pH 7.4).

**Extended Data Fig. 8.**
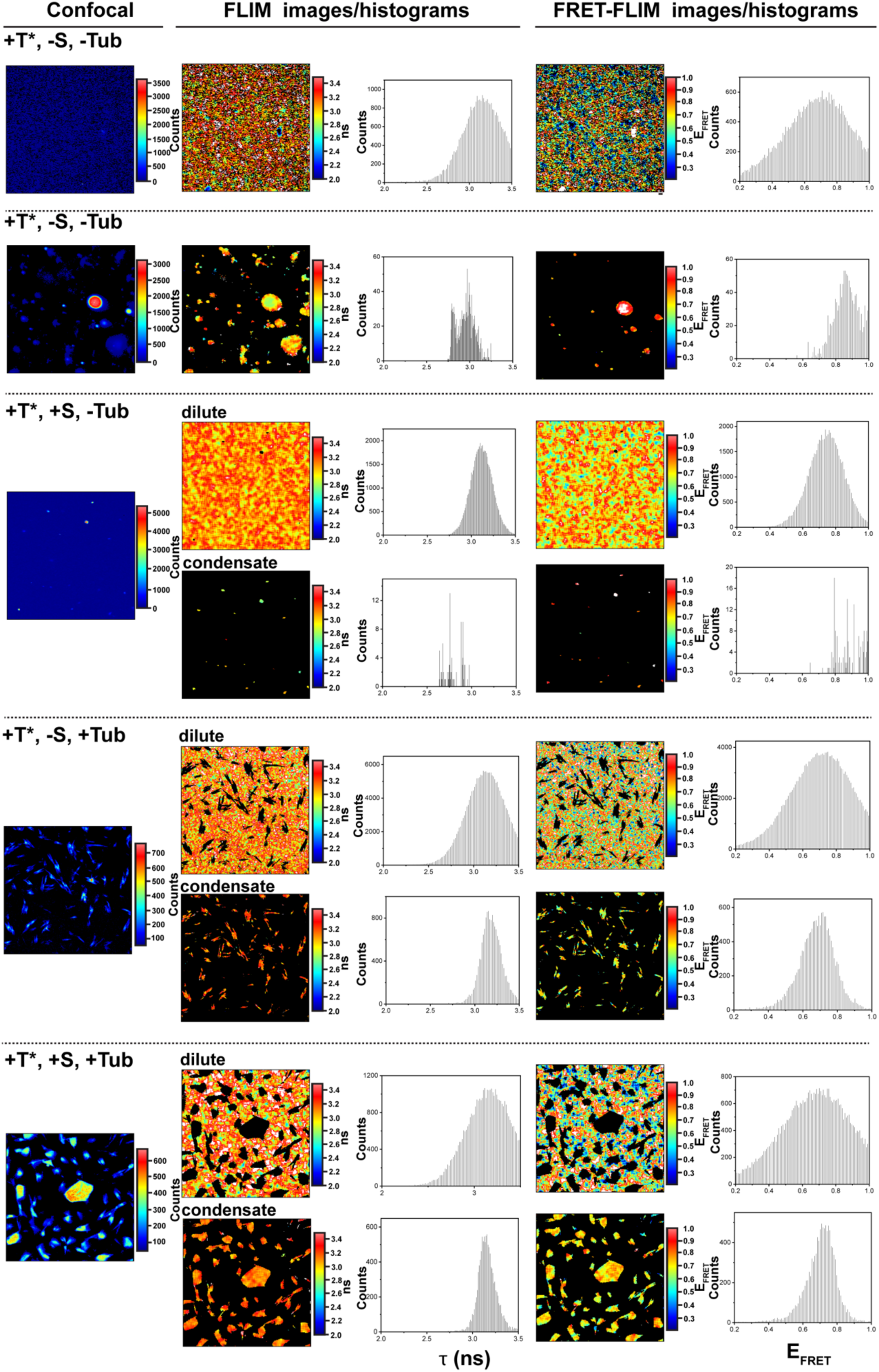
Tau conformational states modulated by αSyn and Tubulin. Representative confocal, FLIM, and FRET-FLIM images with corresponding histograms of fluorescently labeled 1 nM Tau (A488/A594, T*) in the presence 50 µM unlabeled Tau, 50 µM αSyn and/or 10 µM tubulin in FRET-FLIM buffer (5 mM Hepes, 8 mM Pipes, 0.2 mM MgCl_2_, 0.05 mM EGTA, and 1 mM GTP, pH 7.4).

**Extended Data Fig. 9.**
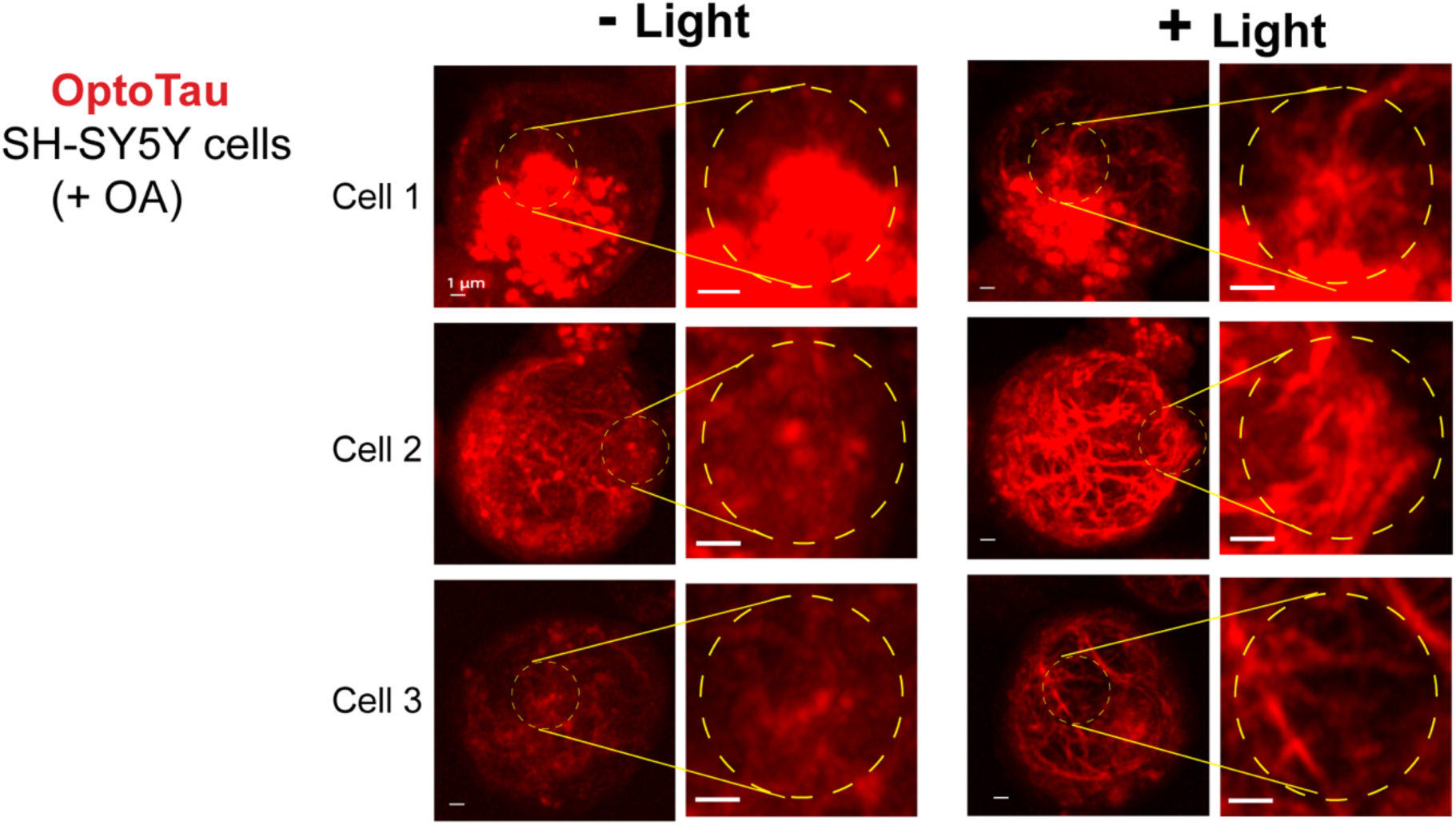
OptoTau enhances microtubule and neurite formation. Confocal images of representative human neuroblastoma SH-SY5Y cells with OptoTau expression (red) with and without light activation (yellow dashed circles). Prior to imaging, the cells were treated with 25 nM okadaic acid for 18 hr to induce cellular stress and microtubule disassembly.

**Extended Data Fig. 10.**
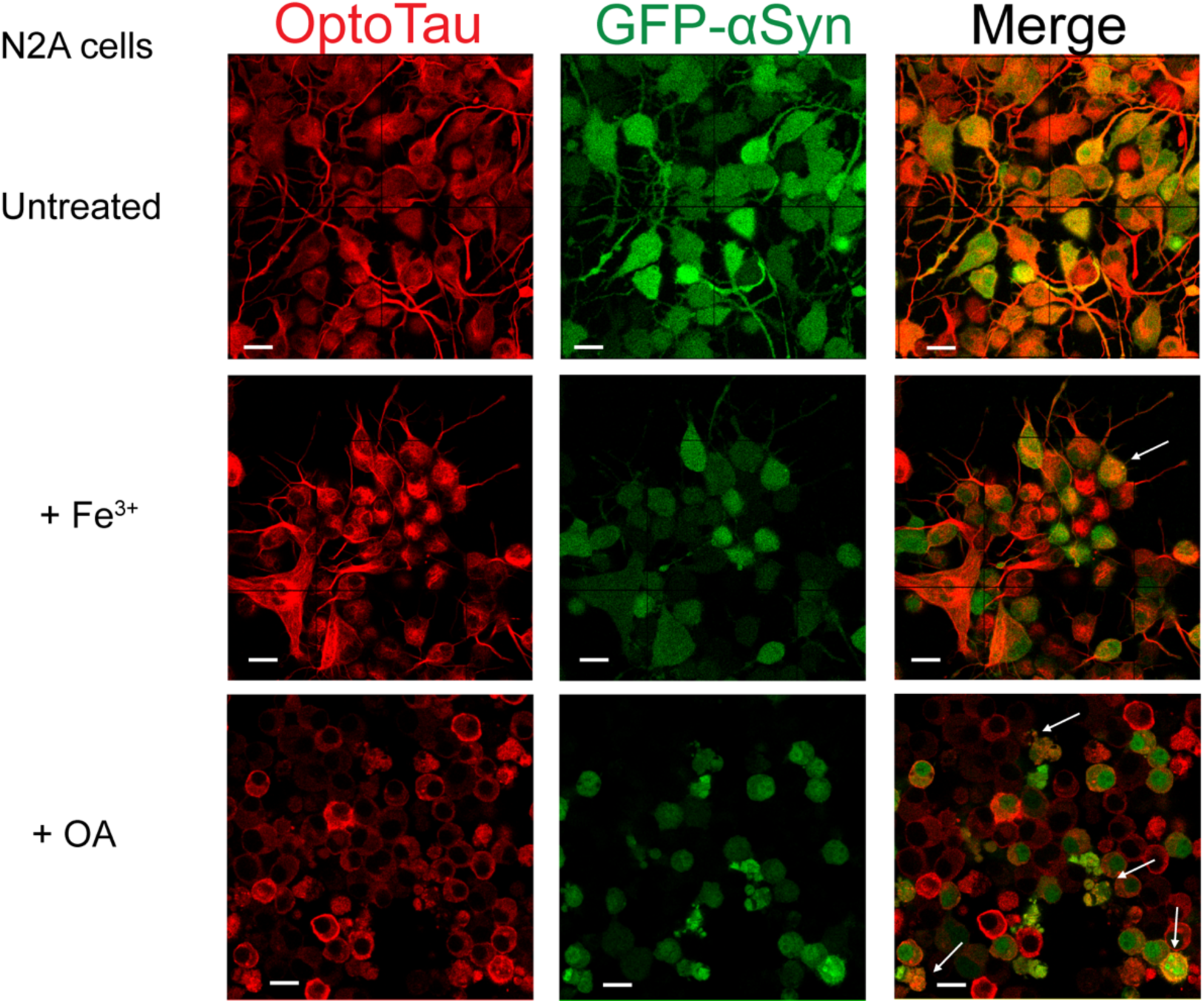
Colocalized Tau and αSyn puncta in stressed N2A cells. Representative confocal images of mouse neuroblastoma N2A cells stably co-expressing OptoTau and GFP-αSyn, treated with 5 mM ammonium iron (III) citrate (Fe^3+^) for 18 hr or 25 nM okadaic acid (OA) for 18 hr. Colocalized Tau/αSyn puncta (yellow) are indicated using white arrows. Scale bar = 20 µm.

**Extended Data Table 1.** FRET-FLIM Sample Protein Sample Compositions.

| Conditions | Tau<sup>A290CE380C</sup> (A488/A594) | Tau<sup>WT</sup> (50 μM) | αSyn<sup>G7CG84C</sup> (A488/A594) | αSyn<sup>WT</sup> (50 μM) | Tubulin (10 μM) | LLPS |
| --- | --- | --- | --- | --- | --- | --- |
| T*S-Tub- | + | - | - | - | - | - |
| T*+S-Tub- | + | + | - | - | - | + |
| T*+S+Tub- | + | + | - | + | - | + |
| T*+S-Tub+ | + | + | - | - | + | + |
| T*+S+Tub+ | + | + | - | + | + | + |
| S*T-Tub- | - | - | + | - | + | - |
| S*T-Tub+ | - | - | + | - | + | - |
| S*T+Tub- | - | + | + | - | - | + |
| S*T+Tub+ | - | + | + | - | + | + |

