## Supplementary Methods for "Tubulin transforms Tau and α-synuclein condensates from pathological to physiological"

**OptoTau (Tau-mCh-Cry2WT) primers.**

Tau-1F 5’-cgg tga tcg tag cgg tta tag cag tcc gg-3’
Tau-1R 5’-ccg gac tgc tat aac cgc tac gat cac cg-3’

Tau-2F 5’-ggt gga gga atg gtg tct aaa g-3’
Tau-2R 5’-ggt agc tcc gga tcc act gtc-3’

**Tau constructs primers.**

C196G forward 5’-cgg tga tcg tag cgg tta tag cag tcc gg-3’
reverse 5’-ccg gac tgc tat aac cgc tac gat cac cg-3’
A291C forward 5’-gag caa cgt tca gag caa atg cgg tag caa aga taa cat taa ac-3’
reverse 5’-gtt taa tgt tat ctt tgc tac cgc att tgc tct gaa cgt tgc tc-3’
S241C forward 5’-gag cag cgc aaa atg ccg tct gca gac cg-3’
reverse 5’-cgg tct gca gac ggc att ttg cgc tgc tc-3’
S341C forward 5’-tgg cca ggt tga agt taa atg cga aaa act gga ttt caa ag-3’
reverse 5’-ctt tga aat cca gtt ttt cgc att taa ctt caa cct ggc ca-3’
E380C forward 5’-cat aaa ctg acc ttt cgc tgc aat gca aaa gcc aaa acc g-3’
reverse 5’-cgg ttt tgg ctt ttg cat tgc agc gaa agg tca gtt tat g-3’
C401G forward 5’-tcc ggt tgt tag cgg tga tac cag tcc gc-3’
reverse 5’-cgc gac tgg tat cac cgc taa caa ccc ga-3’
